# OXPHOS Promotes Apoptotic Resistance and Persistence in T_H_17 cells

**DOI:** 10.1101/2021.10.01.462812

**Authors:** Hanna S. Hong, Nneka E. Mbah, Mengrou Shan, Kristen Loesel, Lin Lin, Peter Sajjakulnukit, Anthony Andren, Atsushi Hayashi, Brian Magnuson, Zhaoheng Li, Yuying Xie, Li Zhang, Yu Leo Lei, Anthony W. Opipari, Rafael J. Argüello, Ilona Kryczek, Nobuhiko Kamada, Weiping Zou, Luigi Franchi, Costas A. Lyssiotis

## Abstract

Apoptotic cell death is a cell-intrinsic, immune tolerance mechanism that regulates the magnitude and resolution of T cell-mediated responses. Evasion of apoptosis is critical for the generation of memory T cells, as well as autoimmune T cells, and knowledge of the mechanisms that enable resistance to apoptosis will provide insight into ways to modulate their activity during protective and pathogenic responses. IL-17-producing CD4 T cells (T_H_17s) are long-lived, memory cells. These features enable their role in host defense, chronic inflammatory disorders, and anti-tumor immunity. A growing number of reports now indicate that T_H_17s in vivo require mitochondrial oxidative phosphorylation (OXPHOS), a metabolic phenotype that is poorly induced in vitro. To elucidate the role of OXPHOS in T_H_17 processes, we developed a system to polarize T_H_17s that metabolically resembled their in vivo counterparts. We discovered that directing T_H_17s to use OXPHOS promotes mitochondrial fitness, glutamine anaplerosis, and an anti-apoptotic phenotype marked by high BCL-XL and low BIM. Through competitive co-transfer experiments and tumor studies, we further revealed how OXPHOS protects T_H_17s from cell death while enhancing their persistence in the periphery and tumor microenvironment. Together, our work demonstrates a non-classical role of metabolism in regulating T_H_17 cell fate and highlights the potential for therapies that target OXPHOS in T_H_17-driven diseases.

## Introduction

Apoptosis regulates T cell development, immune homeostasis, and tolerance (ElTanbouly and Noelle, 2021; Hildeman et al., 2007; Rathmell and Thompson, 2002). Upon pathogen encounter, activated T cells clonally expand, differentiate into effector cells, and produce effector cytokines. Following pathogen clearance, upwards of 95% of effector T cells undergo apoptotic cell death. During this contraction phase, sufficient cell death is critical to prune the immune response and limit unnecessary inflammation. The remaining cells that escape apoptosis persist as memory cells that provide long-term protection against re-infection. Thus, there exists a balance between the cell survival required for immunologic memory and the cell death necessary to prevent excessive tissue damage. The factors that underlie this balance have been extensively studied in CD8 T cells, yet remain incompletely defined in CD4 T cell subsets (e.g., T_H_1, T_H_17, T_reg_).

Glycolysis and mitochondrial oxidative phosphorylation (OXPHOS) are the two principal bioenergetic pathways in a cell, and their differential activity has major impacts on virtually all aspects of cellular metabolism. Predictably, each of these have similarly been implicated in controlling T cell fate and function. More recent studies have further refined our understanding of the metabolic programs that regulate T cells, which have revealed that T cell subsets adopt distinct and often times unique metabolic dependencies to execute subset-specific functions (Geltink et al., 2018). Nevertheless, while the majority of immunometabolism studies have focused on the role of metabolism during T cell differentiation (Berod et al., 2014; Gerriets et al., 2015; Johnson et al., 2018; Roy et al., 2020; Shin et al., 2020; Xu et al., 2017) and memory cell generation and responses (Adams et al., 2016; Buck et al., 2016; Sukumar et al., 2013; van der Windt et al., 2012), it remains elusive how metabolic activity during the effector phase regulates their fate during the contraction phase.

IL-17-producing CD4 T cells (T_H_17s) are a subset of CD4 T cells critical for host-defense and mucosal immunity. T_H_17s are long-lived cells with self-renewing capacity, features that facilitate their role in autoimmunity and, conversely, anti-tumor immunity in the setting of adoptive T cell transfer (Knochelmann et al., 2020; Kryczek et al., 2011; Martin-Orozco et al., 2009; Muranski et al., 2008; Muranski et al., 2011; Shi et al., 2009). Our knowledge of T_H_17 metabolism largely relies on in vitro studies that identified T_H_17s as predominately glycolytic, resulting in the presumption that T_H_17s in vivo would also be glycolytic. However, attempts to inhibit glycolysis in T_H_17s during established disease showed little therapeutic efficacy (Berod et al., 2014; Gerriets et al., 2015). Growing evidence now indicate that in vivo T_H_17s require OXPHOS for lineage specification and pathogenicity (Franchi et al., 2017; Kaufmann et al., 2019; Shin et al., 2020). These findings implicate the role of OXPHOS in T_H_17 effector function, but it remains unknown why T_H_17s require OXPHOS, and how immune responses are modulated by the use of OXPHOS.

In this study, we examined the role of OXPHOS in T_H_17 biology. We developed a culture system amenable to manipulation and capable of retaining features of naturally arising T_H_17s. With this, we found that the predominate use of glycolysis or OXPHOS can comparably support T_H_17 differentiation and function. In contrast, we demonstrate that T_H_17s forced to utilize OXPHOS are provided considerable resistance to apoptotic cell death. We found that this was due to metabolic reprogramming, which results in the upregulation of the anti-apoptotic protein BCL-XL and downregulation of the pro-apoptotic activator BIM. This anti-apoptotic program intrinsic of OXPHOS T_H_17s led to enhanced persistence in the periphery and the tumor microenvironment, suggesting that OXPHOS provides a survival advantage in T_H_17s and supports their long-lived phenotype.

## Results

### OXPHOS promotes metabolic fitness in T_H_17 cells

In vivo, T_H_17 cells require mitochondrial respiration for effector functions (Franchi et al., 2017; Kaufmann et al., 2019; Shin et al., 2020). However, current techniques are limited in their ability to study T_H_17 metabolism in vivo (Voss et al., 2021); rapid ex vivo analysis is bounded by cell number, and T_H_17 expansion in culture leads to changes in both the metabolic and functional phenotype. To overcome these limitations, we developed a cell culture system in which we can maintain T_H_17 cells in their native metabolic state during expansion, allowing for adequate cell numbers for detailed metabolic and molecular analysis.

To this end, naïve CD4 T cells were isolated from the spleen and lymph nodes of mice and activated under T_H_17-polarizing conditions in glucose-free media supplemented with an equimolar concentration of galactose (*i*.*e*. 10mM). As a structural isomer of glucose, galactose inefficiently enters glycolysis through the Leloir pathway, which is thought to impede glycolytic generation of ATP (Bustamante and Pedersen, 1977). In this way, T_H_17 cells are forced to utilize mitochondrial OXPHOS to generate ATP, and thus adopt an oxidative metabolic program mirroring that used by T_H_17 cells in vivo. In parallel, T cells were polarized using the conventional paradigm in standardized media that contained 10mM glucose. We herein refer to cells differentiated in galactose-containing media as OXPHOS T_H_17s and those differentiated in glucose as glycolytic T_H_17s (**Fig. 1A**).

**Figure 1:**
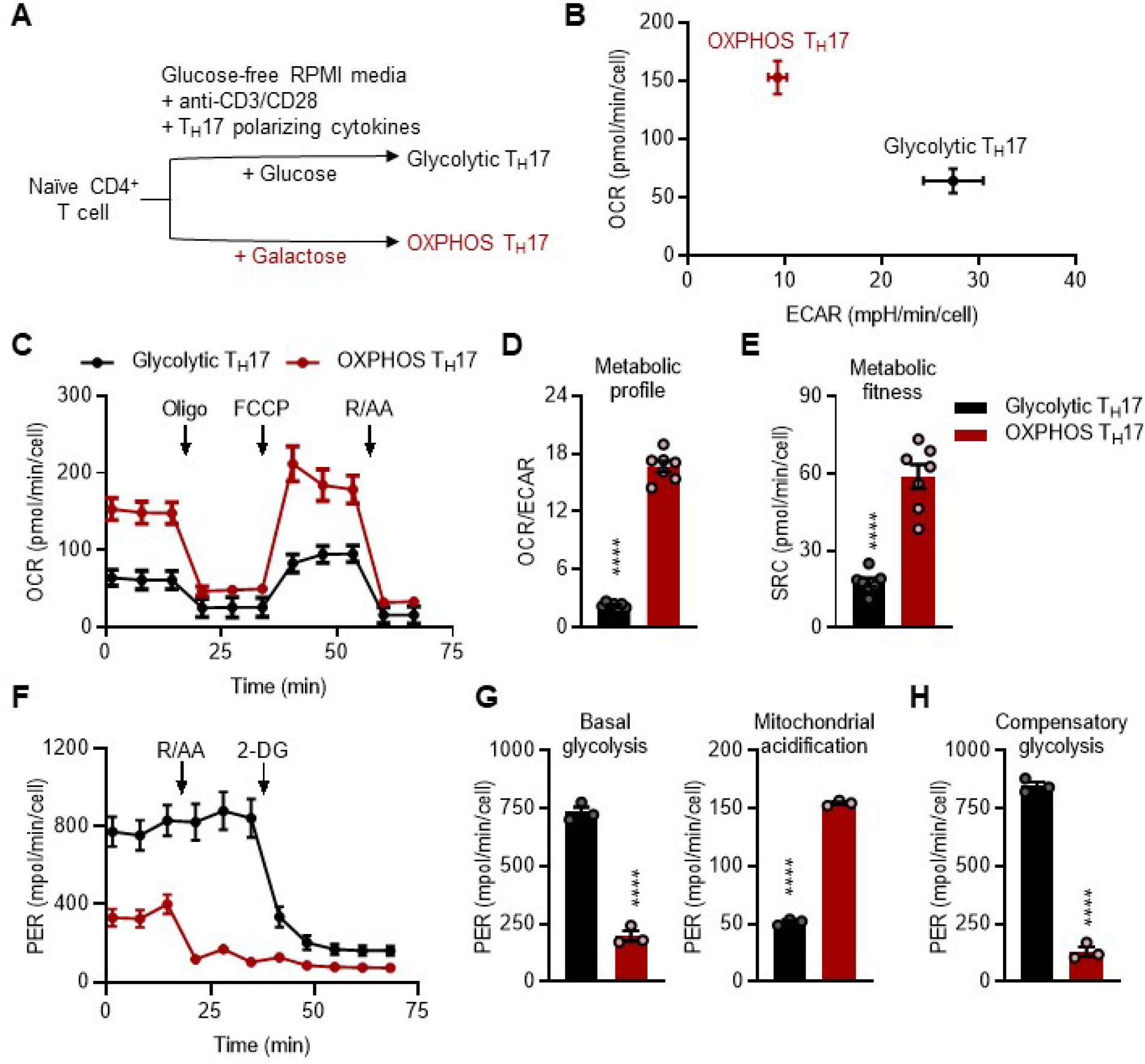
OXPHOS promotes metabolic fitness in T_H_17s. (A) Experimental schema: naïve CD4^+^ T cells were activated with anti-CD3/CD28 and cultured in either glucose- or galactose-containing media in the presence of T_H_17 polarizing cytokines. (B-E) Seahorse flux analysis of glycolytic and OXPHOS T_H_17s using the mitostress test. The metabolic phenotype was determined based on the ratio of OCR/ECAR and metabolic fitness by the spare respiratory capacity (n=6). Graphs represents one of six independent experiments. (F-H) Seahorse flux analysis using the glycolytic rate assay. (F) The total proton efflux rate (PER) reflects media acidification due to glycolytic activity and mitochondrial metabolism. (G) Basal glycolytic activity (left) was determined by subtracting total PER from media acidification produced by mitochondrial metabolism (right). (H) Maximal glycolytic activity (PER values after rotenone and antimycin A co-treatment) reflects compensatory glycolysis (n=3). All data are mean ±SEM. (D,E,G, and H) Statistical significance was determined using a two-tailed unpaired t-test (****p<0.001).

To determine whether our culture system induces distinct metabolic phenotypes, and notably an oxidative phenotype reflective of T_H_17s in vivo (Franchi et al., 2017), we first utilized Seahorse flux analysis to perform the mitochondrial stress test. T_H_17s cultured in galactose-containing media had higher oxygen consumption rates (OCR), lower extracellular acidification rates (ECAR), and an increased ratio of OCR to ECAR, as compared to that of cells cultured in glucose-containing media (**Fig. 1B-D**). Further, OXPHOS T_H_17s exhibited a higher spare respiratory capacity (SRC), which is suggested to be a measurement of the reserved energy in a cell for responses to stress, also referred to as metabolic fitness (**Fig. 1E**). In the context of T cell biology, SRC was demonstrated to be a prerequisite for the generation of memory CD8 T cells that provide superior immunity (van der Windt et al., 2012). Thus, manipulation of media composition can skew metabolic phenotypes in vitro, and OXPHOS T_H_17s exhibit increased metabolic fitness, similar to that of long-lived CD8 T cells.

### Standardized culture conditions promote maximal glycolysis in T_H_17s

Galactose carbon, after phosphorylation and isomerization, enters glycolysis as glucose 6-phosphate, in a manner akin to glucose. Thus, to study the bioenergetic use of glycolysis between our OXPHOS and glycolytic T_H_17s, the glycolysis rate assay was performed. Specifically, we employed the proton efflux rate (PER), which differentiates media acidification produced by glycolytic or mitochondrial activity (**Fig. 1F**). Glycolytic T_H_17s exhibited higher basal glycolysis compared to OXPHOS T_H_17s, and mitochondrial activity was the major contributor to media acidification in OXPHOS T_H_17s (**Fig. 1G**). To assess glycolytic capacity, cells were co-treated with the mitochondrial poisons rotenone and antimycin A to inhibit the electron transport chain (ETC) and promote the use of glycolysis as the main source of energy. OXPHOS T_H_17s were unable to upregulate glycolysis to compensate for ETC inhibition (**Fig. 1H**). In contrast, glycolytic T_H_17s exhibited a modest increase in PER upon ETC inhibition, relative to baseline, suggesting that T_H_17s cultured in standardized media already exhibit near maximal glycolytic activity.

### Aerobic glycolysis is not required for T_H_17 differentiation or effector function

T cell differentiation is primarily driven by immunological cues, such as co-stimulation and the cytokine environment (Murphy and Stockinger, 2010; Yamane and Paul, 2012). A number of recent studies have demonstrated how the metabolic state of a cell can also play a crucial role in regulating T cell identity (Mehta et al., 2017; Zaslona and O’Neill, 2020). To examine whether the predominate use of OXPHOS alters T_H_17 differentiation, cells were intracellularly stained for the T_H_17 lineage-defining transcription factor RORγt (**Fig. 2A**). Since T_H_17s have been observed to transdifferentiate under certain circumstances in vivo, the expression of Tbet or Foxp3 were also examined to assess whether OXPHOS induces T_H_1 or T_reg_ transdifferentiation, respectively. Relative to T_H_1 or T_reg_ controls, T_H_17s retained their T_H_17 identity, irrespective of metabolic phenotype (**Fig. 2B, 2C**). Further, glycolytic and OXPHOS T_H_17s expressed their effector cytokine IL-17 at comparable levels, a reflection of expression of the lineage-defining transcription factor RORγt, and were low-producers of IFNγ (**Fig. 2D, 2E**).

**Figure 2:**
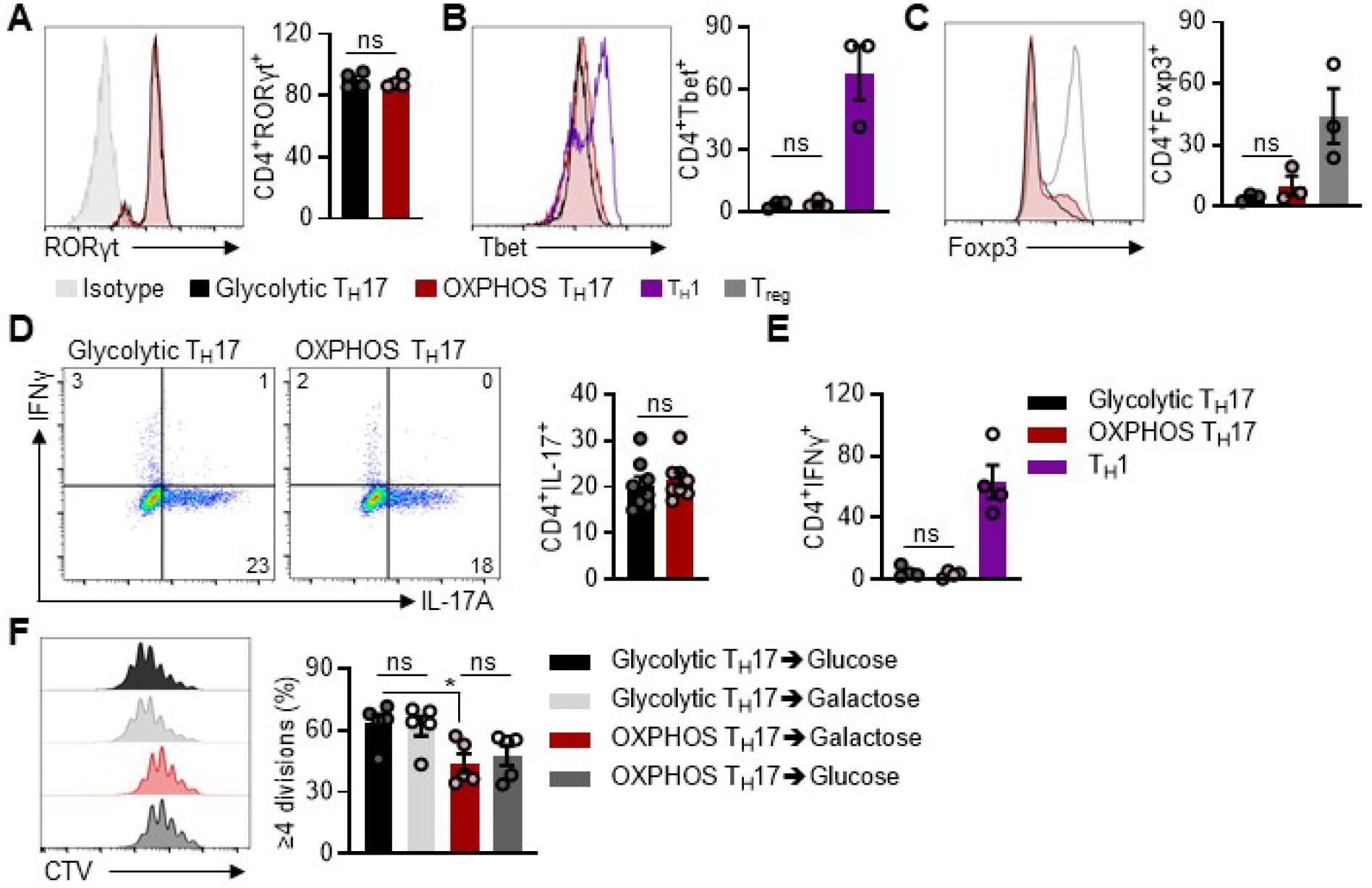
Aerobic glycolysis is not required for T_H_17 differentiation or function. (A) Representative flow histogram of RORγt expression (left) and accompanying quantification (right) expressed as the percentage of RORγt^+^ cells (n=4). (B-C) Quantification of Tbet or Foxp3 expression in glycolytic and OXPHOS T_H_17s, as compared to T_H_1 or T_reg_ controls, respectively (n=3). (D-E) Effector cytokine profile of T_H_17s (D) Comparison of IL-17A^+^ expression between T_H_17s (n=8), or (E) IFNγ^+^ expression in T_H_17s, relative to T_H_1 controls (n=3). (F) Cell Trace Violet (CTV)-stained T_H_17s were polarized under glycolytic or OXPHOS conditions for 5 days, and then split into either glucose- or galactose-containing media for 48hrs. Representative flow histogram (left) and accompanying quantification (right) of T_H_17 cell proliferation (n=4). Expression of intracellular markers and stains were live-gated on CD4^+^ single cells. All results represent at least 3 independent experiments. All data are mean ±SEM. (A-F) Statistical significance was determined using one-way ANOVA (*p<0.05).

Cell proliferation is intimately linked to metabolic activity, as glycolysis and OXPHOS produce the energy and biosynthetic intermediates needed for cell growth. To assess proliferation, Cell Trace Violet (CTV)-dilution experiments were performed and revealed that OXPHOS T_H_17s proliferated at a slower rate compared to that of glycolytic T_H_17s (**Fig. 2F**). To examine whether the proliferative phenotype induced by the metabolic culture conditions is fixed, OXPHOS T_H_17s were subcultured in glucose-containing media in an effort to promote the use of glycolysis. We found that the decreased proliferative rate of OXPHOS T_H_17s persisted following subculture in glucose-containing media for 48hrs; likewise, the proliferative rate of glycolytic T_H_17s was unperturbed when pressured to use OXPHOS through subculture in galactose-containing media. Based on these collective data, we infer that a stable metabolic program was established in T_H_17s based on metabolic substrates applied during polarization.

### Profiling analyses reveal differential glutamine metabolism between glycolytic and OXPHOS T_H_17s

Our data indicate that differential utilization of glycolysis or OXPHOS does not impact T_H_17 proliferation, differentiation, or cytokine production. This flexibility in metabolic programs during differentiation allowed for the examination into *non*-energetic functions of glycolysis and OXPHOS in an otherwise homogenous T_H_17 population. In addition, this strategy also allowed for a side-by-side assessment of the role of glycolysis vs. OXPHOS within the T_H_17 cell type.

To this end, we profiled T_H_17 cultures by bulk RNA sequencing (RNA-seq) and liquid chromatography-coupled tandem mass spectrometry (LC-MS/MS)-based metabolomics. First, RNAseq analysis revealed that nearly 17% of detected genes were differentially expressed (**Fig. 3A**). Of the top 500 most significantly changed genes, Gene Ontology analysis revealed enrichment in processes involved in immune differentiation and cytokine production, metabolism, regulatory activity, and cell death (**Fig. 3B**). Gene set enrichment analysis identified pathways associated with various signaling events, hypoxia, inflammation, and apoptosis in glycolytic T_H_17s (**Fig. S1A**). Examination of genes required for ETC activity, as well as some involved in glycolysis, revealed that these were upregulated in OXPHOS T_H_17s (**Fig. S1B**).

**Figure 3:**
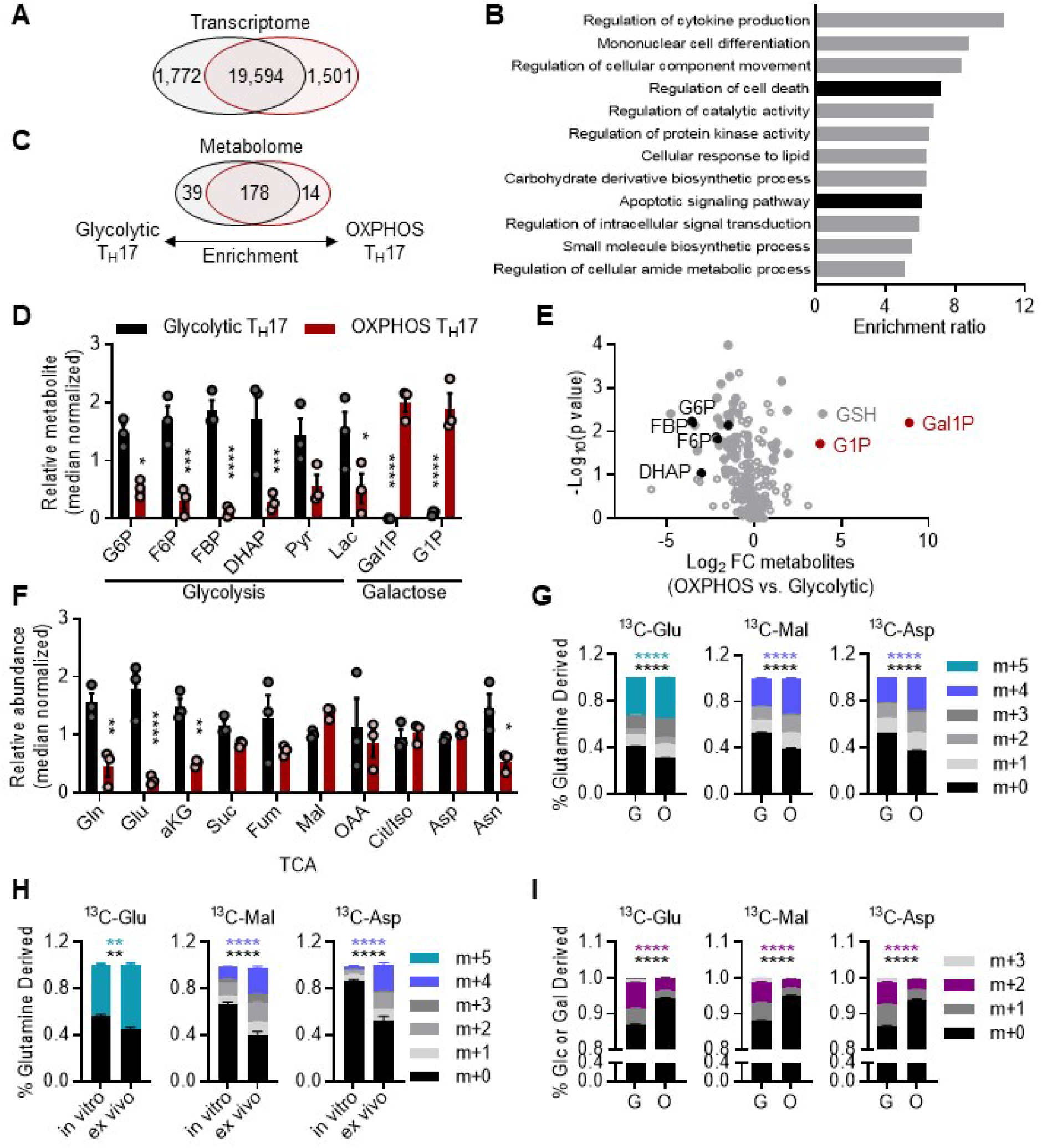
Transcriptomic and metabolomic profiling of OXPHOS and glycolytic T_H_17s. (A) Venn diagram depicting the number of differentially expressed genes (fold change>|0.5|, p-value<0.05) between glycolytic (black) and OXPHOS (red) T_H_17s analyzed by RNA-seq. (B) Gene Ontology analysis of the top 500 most significantly changed genes from (A). (C) Venn diagram depicting the number of differential metabolites (fold change>|1|, p-value<0.1) between glycolytic and OXPHOS T_H_17s analyzed by metabolomics. (D) Differential metabolite levels of intermediates in glycolysis or galactose metabolism in T_H_17s. (E) Volcano plot of metabolites based on fold change (FC) and corresponding significance. Each circle represents a metabolite, significantly changed metabolites (log_2_FC >|1|, p-value<0.1) are depicted as solid circles. (F) Differential metabolite levels of TCA intermediates in T_H_17s. (G and H) [U]^13^C glutamine tracing metabolomics in (E) glycolytic vs. OXPHOS T_H_17s or (F) in vitro vs. ex vivo T_H_17 cells. Fractional labeling pattern of isotopologues for TCA metabolites (mass+0-5; m+): ^13^C glutamine-derived glutamate (^13^C-Glu; m+5), malate (^13^C-Mal; m+4), and aspartate (^13^C-Asp; m+4). (I) [U]^13^C glucose incorporation in glycolytic T_H_17s was compared to [U]^13^C galactose incorporation in OXPHOS T_H_17s. Fractional labeling pattern of TCA metabolites (m+2). All data are mean ±SEM (n=2-3; biological replicates). Statistical analysis was determined by (D, F, G-I) two-way ANOVA or (E) two-tailed unpaired t-test (*p<0.05; **p<0.01; ***p<0.001; ****p<0.0001). Glucose-6-phosphate (G6P), fructose-6-phosphate (F6P), fructose-1,6-bisphosphate (FBP), dihydroxyacetone phosphate (DHAP), pyruvate (Pyr), lactate (Lac), galactose-1-phosphate (Gal1P), glucose-1-phosphate (G1P), reduced glutathione (GSH), glutamine (Gln), glutamate (Glu), αketoglutarate (aKG), succinate (Suc), fumarate (Fum), malate (Mal), oxaloacetate (OAA), citrate/isocitrate (Cit/Iso), aspartate (Asp), asparagine (Asn).

As metabolic pathway activity is influenced by numerous factors (e.g., nutrient uptake, metabolic gene and protein expression) and culminates in the production of metabolites, we next examined our metabolomics dataset to compare the metabolite profiles between glycolytic and OXPHOS T_H_17s. Approximately 23% of central carbon metabolites were differentially enriched between T_H_17 types (**Fig. 3C**). In agreement with our Seahorse flux analysis, we found that the intermediates of glycolysis were more abundant in glycolytic T_H_17s, while intermediates of galactose metabolism were abundant in OXPHOS T_H_17s (**Fig. 3D, 3E**). These data reproduce our previous metabolomics analysis on in vitro-vs. in vivo-derived T_H_17 cells (Franchi et al., 2017), wherein the in vivo cells have much smaller glycolytic pools, further substantiating the culture system being explored herein. In fact, carbohydrate metabolism was among the most distinguishing features according to metabolic pathway analysis (**Fig. S1C**).

Despite setting the carbohydrate content in our T_H_17 cultures, glutamine metabolism and related pathways (e.g., glutathione, non-essential amino acid metabolism, pyrimidine metabolism) were even more enriched than carbohydrate metabolism-related pathways (**Fig. S1C-E**). Indeed, recent studies have illustrated that T_H_17s rely on glutamine metabolism to support differentiation. Glutamine serves as one of the major carbon sources for the tricarboxylic acid (TCA) cycle, the products of which mediate T_H_17 lineage specification (Xu et al., 2017). Glutamate, the product of glutamine deamidation, is also required for the synthesis of reduced glutathione (GSH), which is critical in T_H_17s to limit redox stress during differentiation (Johnson et al., 2018; Mak et al., 2017). While we found that glycolytic and OXPHOS cultures supported T_H_17 differentiation equivalently (**Fig. 2**), our metabolomics analysis indicated that glutamine, glutamate, and αKG were depleted in OXPHOS T_H_17s, with similar trends for most of the other TCA cycle intermediates, as compared to those in glycolytic T_H_17s (**Fig. 3F**). Furthermore, GSH was among the most differentially regulated metabolites, being enriched >10-fold in the OXPHOS cultures (**Fig. 3E**).

Thus, to compare how glycolytic and OXPHOS T_H_17s use glutamine, we performed stable isotope tracing metabolomics by culturing T_H_17s in the presence of uniformly 13-carbon-labeled ([U]^13^C)-glutamine. From this analysis, we found that OXPHOS cultures increased the incorporation of glutamine-derived carbon into TCA cycle intermediates and de novo GSH biosynthesis (**Fig. 3G, S2A**), suggesting that they have a higher rate of both glutamine consumption and metabolism. To corroborate these findings, we compared [U]^13^C-glutamine labeling patterns in T_H_17s derived under glycolytic conditions in vitro to those derived in vivo and then cultured/labeled ex vivo. Here again, we see a clear enrichment of glutamine-derived carbon in the TCA cycle intermediates from the oxidative in vivo cells, relative to the in vitro glycolytic cells (**Fig. 3H, S2B**).

In parallel with the glutamine tracing studies above, we also traced carbohydrate metabolism in our T_H_17 cultures using [U]^13^C-glucose for the glycolytic T_H_17s and [U]^13^C-galactose for the OXPHOS T_H_17s. The results from this experiment revealed that the glycolytic cells incorporate more carbohydrate-derived carbon into the TCA cycle than do the OXPHOS cultures (**Fig. 3I**). Together with our data above, this suggests that the OXPHOS cells require more glutamine-derived carbon to fill the TCA cycle to account for the limited entry of galactose carbon.

### OXPHOS T_H_17s are resistant to apoptotic cell death

As indicated above, the predominate use of glycolysis or OXPHOS has widespread impacts on central carbon metabolism. One of the most significantly altered metabolites between glycolytic and OXPHOS T_H_17s was GSH (**Fig. 3D, 4A**). GSH is an antioxidant that detoxifies reactive oxygen species (ROS) through the conversion of reduced GSH to oxidized glutathione (GSSG). Comparison of the ratio of reduced to oxidized glutathione and the levels of mitochondrial and intracellular ROS revealed that OXPHOS T_H_17s had increased antioxidant capacity (**Fig. 4B-D**).

**Figure 4:**
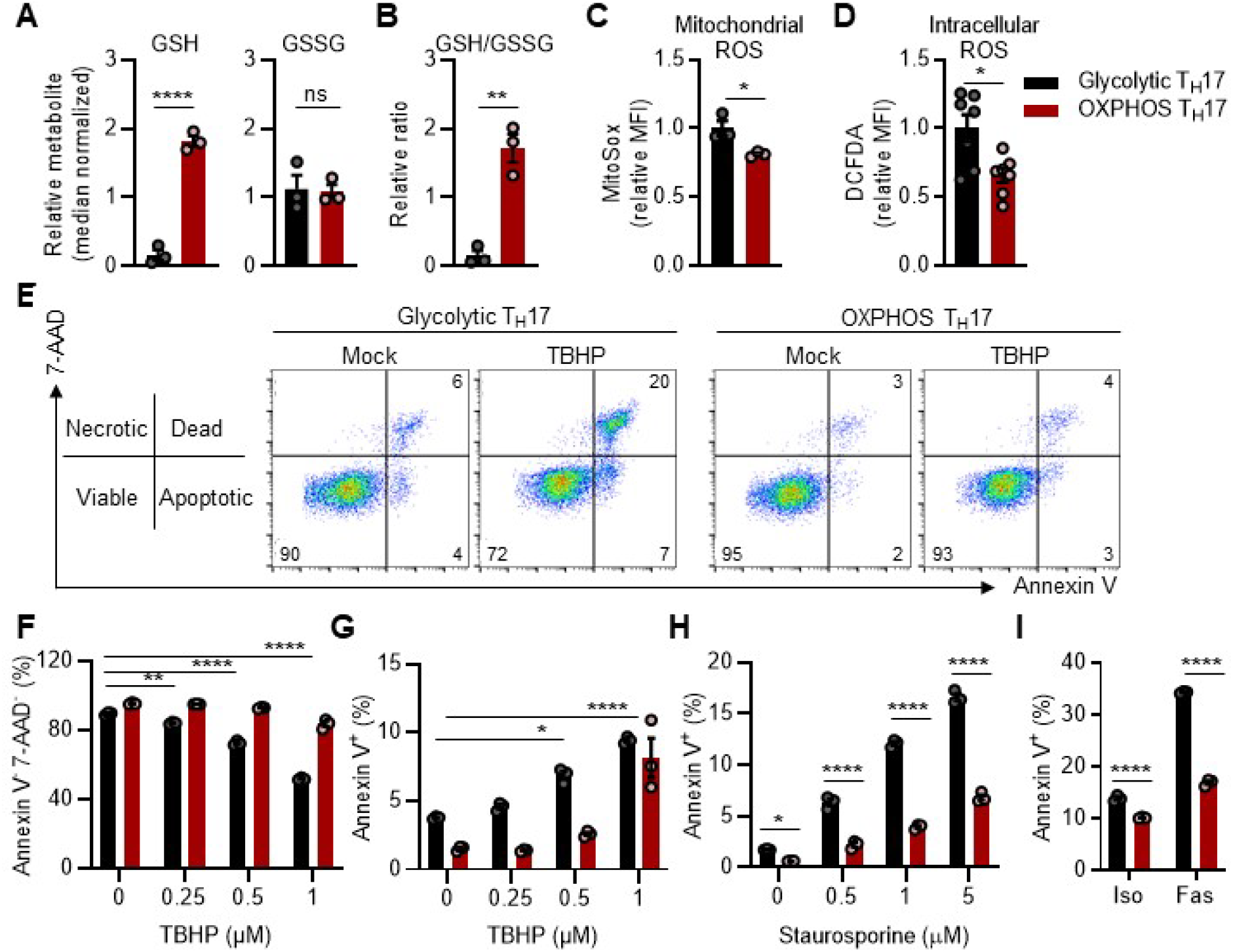
Glycolysis sensitizes T_H_17s to apoptosis. (A-B) Relative abundance of reduced glutathione (GSH) and oxidized glutathione (GSSG) were measured by LC-MS/MS and used to determine the relative ratio of GSH to GSSG in glycolytic and OXPHOS T_H_17s (n=3; biological replicates). (C-D) Assessment of reactive oxygen species (ROS) accumulation in T_H_17s. (C) Mitochondrial and (D) intracellular ROS were detected using MitoSox and H_2_DCFDA, respectively, and median fluorescent intensity (MFI) was normalized to glycolytic T_H_17s (n=3). Data represent one of two independent experiments. (E-G) Examination of apoptotic sensitivity in T_H_17s treated with tert-butyl H_2_O_2_ (TBHP). (E) Representative flow plots of Annexin V vs. 7-AAD staining of T_H_17s following treatment with 0.5μM TBHP. Accompanying quantification of the frequency of (F) viable or (G) apoptotic cells (n=3). (H and I) Quantification of apoptosis (AnnexinV^+^ 7-AAD^−^ cells) following treatment with (H) staurosporine or (I) anti-Fas (n=3). (E-I) Data represent one of six independent experiments. All data are mean ±SEM. Statistical significance was determined by (A-D) two-tailed unpaired t-test or (F-I) two-way ANOVA (*p<0.05; **p<0.01; ****p<0.0001).

To test whether this baseline difference in antioxidant profile has a functional role, T_H_17s were treated with cell-permeable hydrogen peroxide (tert-butyl hydrogen peroxide; TBHP) as an oxidative challenge. Through Annexin V and 7-AAD dye exclusion, quantification of cell viability revealed that glycolytic T_H_17s were more susceptible to oxidant-induced cell death (**Fig. 4E, 4F**).

These results prompted us to investigate whether differences in metabolic activity in T_H_17s differentially sensitized cells to other pathways that cause cell death. Several lines of evidence led us to focus on apoptosis. First, apoptosis is an immune tolerance mechanism that eliminates T cells in nearly all phases of their life span. T_H_17s have been observed to be long-lived cells (Amezcua Vesely et al., 2019; Kryczek et al., 2011; Muranski et al., 2011), indicating that they escape numerous tolerance mechanisms that would result in apoptotic cell death. Second, our transcriptomics analysis revealed differences in apoptotic signaling between glycolytic and OXPHOS T_H_17s, linking metabolism to T_H_17 apoptotic sensitivity (**Fig. 3B, S1A**). Third, analysis of the frequency of apoptotic cells (i.e., Annexin V^+^7-AAD^−^ cells) following treatment with TBHP identified that glycolytic cells underwent apoptosis to a greater extent than their OXPHOS counterparts (**Fig. 4G**). Thus, to determine if glycolytic and OXPHOS T_H_17s were indeed differentially sensitized to apoptosis, cells were treated with staurosporine to induce the intrinsic apoptotic pathway or Fas to activate the extrinsic pathway. The frequency of apoptosis was induced significantly more in glycolytic cells than OXPHOS T_H_17s (**Fig. 4H, 4I**). Together, these data revealed that OXPHOS T_H_17s were more resistant to apoptotic cell death.

### OXPHOS alters the expression of anti- and pro-apoptotic proteins

The mechanisms by which T_H_17 apoptosis is regulated by metabolic programs in vivo remains unclear, due in part to the limited availability of in vivo methods to study metabolism. Accordingly, we applied our in vitro system to examine the mechanism by which OXPHOS imparts resistance to cell death. First, we observed that glycolytic and OXPHOS T_H_17s differed significantly in cell viability, both at baseline (see vehicle-treated controls) and following induction of either intrinsic or extrinsic apoptosis (**Fig. 4F-I**). This indicated that the predominant use of glycolysis and OXPHOS may induce different apoptotic thresholds, which prime T_H_17s to respond accordingly, irrespective of which apoptotic pathway is activated. Consequently, we queried our RNA-seq dataset for key apoptotic genes and found that glycolytic T_H_17s predominately expressed pro-apoptotic genes (**Fig. S3A**).

Next, we compared the levels of the BCL-2 family proteins to examine the balance of pro-vs. anti-apoptotic factors, which dictates apoptotic sensitivity. Consistent with the observed differences in cell viability, OXPHOS T_H_17s expressed higher levels of the anti-apoptotic regulators BCL-XL and MCL-1, and lower levels of the pro-apoptotic activator BIM, compared to glycolytic T_H_17s. (**Fig. 5A-D**). We also observed that cleaved caspase 3 (CC3) was higher in resting glycolytic T_H_17s, yet concurrent analysis showed that these cells were >90% viable (**Fig. 5D, 5E**). This was further confirmed by the lack of PARP cleavage, which is a product of irreversible apoptotic cell death. These findings are in agreement with the notion that caspase 3 is activated upon T cell activation and serves as a self-limiting mechanism that predestines effector T cells for apoptotic cell death (Garrod et al., 2012). Thus, these observations indicated that glycolytic T_H_17s are molecularly primed to die. By contrast, OXPHOS T_H_17s were intrinsically more resistant to apoptosis through downregulated expression of BIM and upregulated expression of BCL-XL and MCL-1.

**Figure 5:**
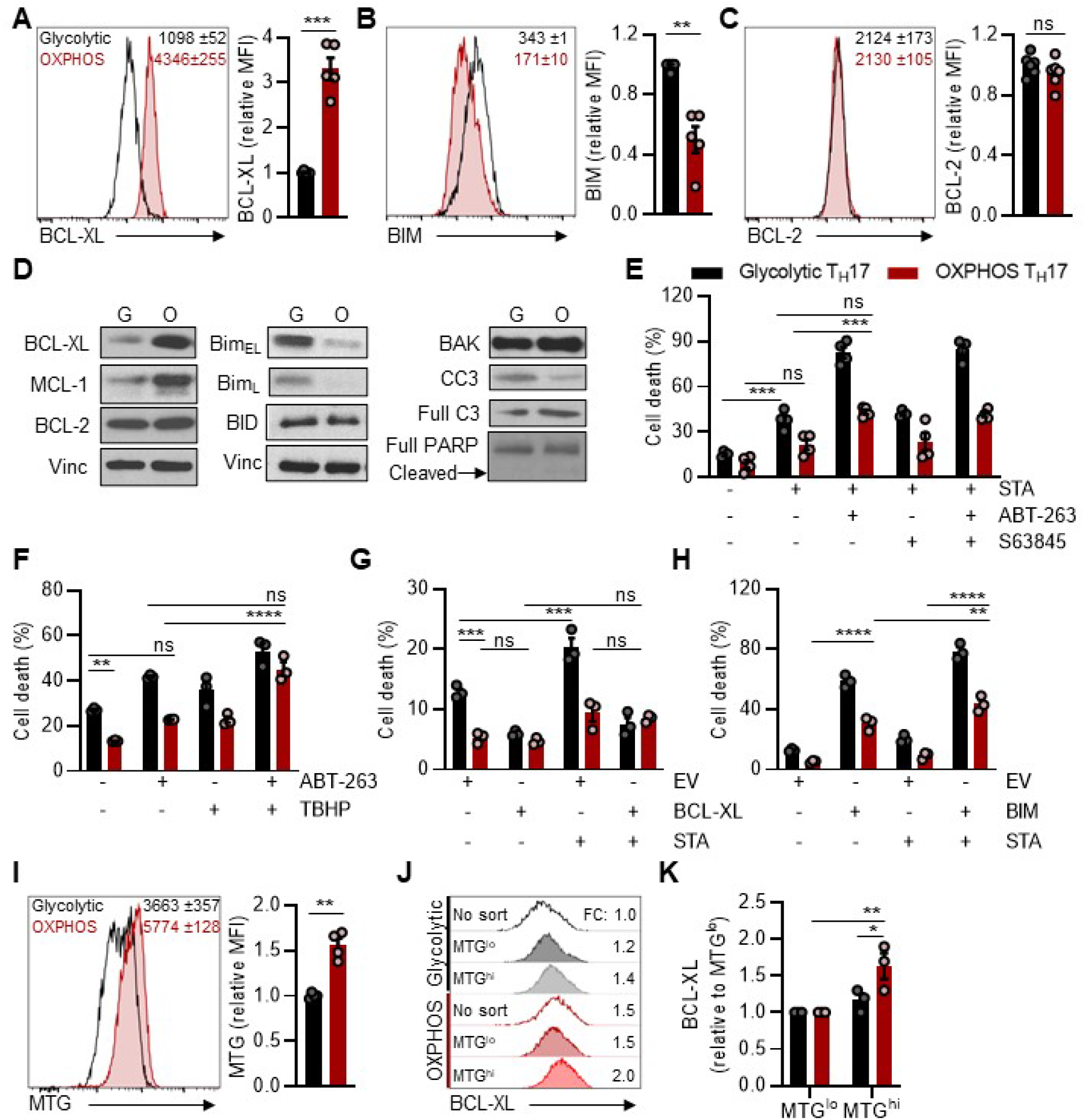
OXPHOS modulates BCL-XL and BIM expression in T_H_17s. (A-D) Expression of anti- and pro-apoptotic regulators in T_H_17s measured by (A-C) flow cytometry or (D) western blotting. Data represent five (A, B; n=5) or two (C; n=6) independent experiments. (E) T_H_17s were treated with ABT-263 and/or S63845, followed by staurosporine (STA). Frequency of cell death was based on Annexin V^+^ 7-AAD^+^ detection by flow cytometry (n=4). Data represent two independent experiments. (F) Frequency of cell death in T_H_17s treated with ABT-263, followed by TBHP (n=3). Data represent one of two independent experiments. (G-H) T_H_17s transduced with retrovirus expressing empty vector (EV) control, (G) BCL-XL or (H) BIM were challenged with STA and cell death was quantified in transduced cells (n=3). Data represent one of two independent experiments. (I) T_H_17s were stained with MitoTracker Green (MTG) to assess mitochondrial mass (n=4). Data represent 4 independent experiments. (J-K) MTG-stained T_H_17s were flow sorted based on the 20% lowest and highest MFI, and then stained for BCL-XL. (J) Flow histogram represents bulk cultures (no sort), MTG^lo^ and MTG^hi^ sorted T_H_17s. Fold change in BCL-XL expression relative to (J) bulk or (K) MTG^lo^ glycolytic T_H_17 cultures (n=3). Data represent 3 independent experiments. All data are mean ±SEM. (A-C, I) Representative flow histograms (left) and quantification of relative MFI (right) in live-gated CD4^+^ T_H_17s and numbers in histograms represent the absolute MFI (mean ±SEM) of 3 technical replicates. Statistical significance was determined by (A-C, I) two-tailed paired t-test or (E-H, K) two-way ANOVA (*p<0.05; **p<0.01; ***p<0.001;****p<0.0001).

As OXPHOS is the relevant metabolic state of T_H_17s in vivo (Franchi et al., 2017), we next assessed whether the anti-apoptotic profile of OXPHOS T_H_17s generated in vitro matched that of in vivo T_H_17 cells. Indeed, analysis of publicly available transcriptome data from T_H_17s generated using comparative in vivo vs. in vitro models revealed that in vivo T_H_17s exhibit an anti-apoptotic gene expression profile, compared to that of in vitro (glycolytic) T_H_17s (**Fig. S3B**).

### BCL-XL is required to mediate apoptotic resistance in T_H_17s

Considering OXPHOS T_H_17s predominately upregulate BCL-XL and MCL-1, we next tested whether apoptotic resistance in OXPHOS T_H_17s is mediated through their anti-apoptotic activity. To this end, T_H_17s were treated with ABT-263, which is a pan-inhibitor of BCL-2, BCL-XL, and BCL-W; S63845 to selectively inhibit MCL-1; or the combination. ABT-263 treatment alone only modestly increased spontaneous apoptosis, presumably due to the low BIM expression inherent to OXPHOS T_H_17s (**Fig. S3C**). However, this modest induction of cell death mediated by ABT-263 treatment brought OXPHOS T_H_17s to the same level of cell death as vehicle-treated glycolytic cells. In contrast, ABT-263 treatment enhanced spontaneous apoptosis in glycolytic T_H_17s, underscoring the importance of BCL-XL, BCL-2, and BCL-W in T_H_17 survival. Treatment with S63845 neither had a significant effect as a single agent nor potentiated the effects of ABT-263 in either glycolytic or OXPHOS T_H_17s (**Fig. S3C**). These results suggest that MCL-1 is not a critical regulator of T_H_17 cell survival.

We next assessed the impact of these anti-apoptotic proteins in protection from instructed apoptotic cell death using the pharmacological inhibitors noted above. T_H_17s were pretreated individually or in combination with ABT-263 and S63845, and then challenged with the apoptotic stimuli staurosporine (STA) or TBHP. ABT-263 potentiated the effects of STA in both glycolytic and OXPHOS T_H_17s, and S63845 again had no impact (**Fig. 5E**). These results further underscore the roles of the BCL-2 family members in T_H_17 apoptosis. In contrast to the results with STA, ABT-263 treatment sensitized OXPHOS T_H_17s to the cytotoxic effects of TBHP to the same degree as glycolytic cultures, suggesting a differential role for BCL-2/BCL-XL in TBHP-mediated apoptosis (**Fig. 5F**).

Based on these results, we then sought to determine if BCL-XL is sufficient to protect T_H_17s from cell death. T_H_17s were retrovirally transduced with empty vector or BCL-XL and treated with STA. BCL-XL expression rescued spontaneous and STA-induced cell death in glycolytic T_H_17s, while having no significant impact on OXPHOS T_H_17s (**Fig. 5G**). Conversely, retroviral-mediated expression of BIM sensitized both OXPHOS and glycolytic T_H_17s to spontaneous and STA-induced cell death (**Fig. 5H**). Together, these data show that the OXPHOS-mediated induction of BCL-XL and repression of BIM facilitate survival and protection from apoptotic cell death, relative to glycolytic controls.

### OXPHOS culture enriches for mitochondrial content and BCL-XL

Mitochondrial integrity is central to cell survival. In naïve cells, T cell activation induces mitochondrial biogenesis and activity to support their exit from cellular quiescence (Ron-Harel et al., 2016; Sena et al., 2013), and manipulation of mitochondrial dynamics in activated CD8 T cells can increase their functionality and persistence in vivo (Buck et al., 2016; van der Windt et al., 2012). With these concepts in mind, we next examined whether differences in mitochondrial dynamics between glycolytic and OXPHOS T_H_17s are responsible for the observed differences in cell survival. Quantification of mitochondrial mass using MitoTracker Green (MTG) indicated that OXPHOS T_H_17s had more mitochondria than glycolytic cultures (**Fig. 5I**). Examination of the regulators of mitochondrial dynamics revealed that OXPHOS T_H_17s upregulated the mitochondrial biogenesis transcription factor PGC1α, relative to glycolytic cultures (**Fig. S4A**). Further, glycolytic T_H_17s exhibited greater phosphorylation of Dynamin-related Protein 1 (DRP1), which drives mitochondrial fission, suggesting that mitochondria in glycolytic cells may undergo more division and subsequent elimination (**Fig. S4B**). Changes in mitochondrial fusion marker OPA1 (Dynamin-like 120 kDa protein) were similar between T_H_17 cultures.

BCL-XL is expressed in the mitochondria, and we hypothesized that the greater mitochondrial content in OXPHOS cultures was responsible for its elevated expression. To test this hypothesis, T_H_17s were stained with MitoTracker Green, the top and bottom 20% of stained cells were sorted into mitochondrial high and low fractions, and then stained for BCL-XL and BIM. OXPHOS T_H_17s with more mitochondria expressed the highest levels of BCL-XL (**Fig. 5J, 5K**). We also observed a modest but significant increase in BIM expression in the MTG high cultures, which was comparable between the glycolytic and OXPHOS cultures (**Fig. S4C**). Together, these results indicated that T_H_17s that predominately use OXPHOS have greater mitochondrial mass, correlating with more BCL-XL.

### OXPHOS promotes T_H_17 persistence in vivo

In the periphery, T cell numbers remain relatively constant in the absence of infection (Jameson, 2002; Surh and Sprent, 2008). T cell survival is mediated by sustained expression of anti-apoptotic BCL-2 family proteins and survival signals. Apoptosis executes the deletion of T cells that fail to compete for limited resources, by way of maintaining immune homeostasis. To examine whether the use of glycolysis vs. OXPHOS in T_H_17s confers differential cell survival and persistence in vivo, we performed competitive co-transfer experiments. First, congenically distinct glycolytic and OXPHOS T_H_17s were generated in culture, mixed in equal proportion (1:1), and adoptively transferred into lymphopenic recipient mice. Recovery of transferred cells 28 days post-transfer revealed that a greater frequency of cells derived from the OXPHOS fraction persisted in the blood, peripheral lymph nodes (pLN), spleen, and lamina propria (LP) (**Fig. 6A**). Importantly, viability analysis of recovered cells indicated that cells from the glycolytic fraction underwent more cell death (**Fig. 6B**). Together, these data indicate that OXPHOS polarization conditions impart greater persistence of T_H_17s in vivo.

**Fig 6:**
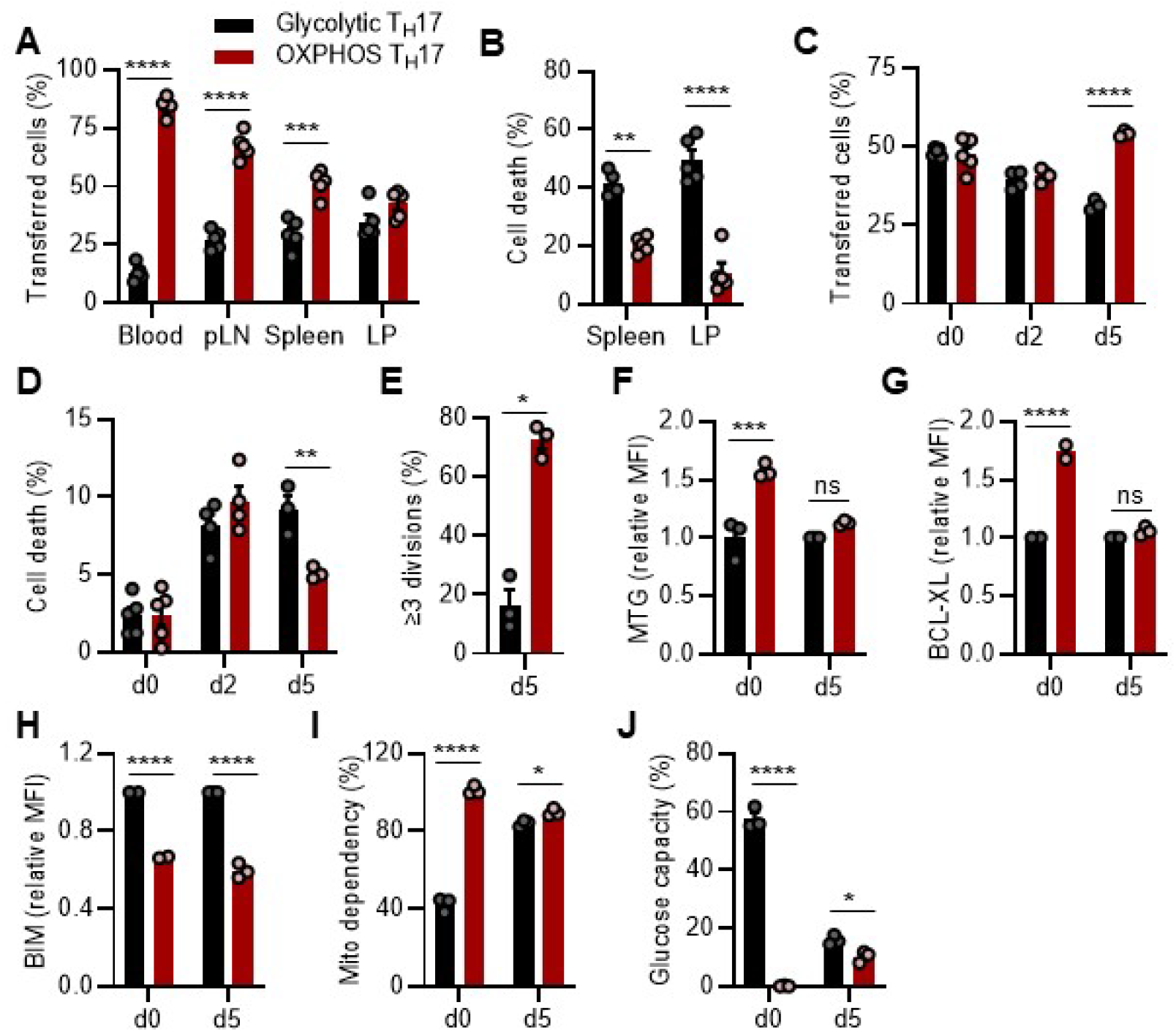
OXPHOS promotes T_H_17 survival and persistence in vivo. (A and B) T_H_17s cultured under glycolytic (CD45.2^+^) or OXPHOS (CD45.1^+^) conditions were equally mixed, and transferred into lymphodeplete recipient mice. After 28 days, recovery of transferred cells was determined based on the frequency of CD45.2^+^ vs. CD45.1^+^ T cells (n=5 mice). (C-E) Glycolytic (CD45.2^+^) or OXPHOS (CD45.1^+^) T_H_17s were CTV labeled, equally mixed, then transferred into C57Bl/6 recipient mice. (C) Cell survival, (D) death, or (E) proliferation was assessed in CTV^+^-gated cells and transferred cells were discriminated based on congenic markers (n=3-4 mice). (F-J) CTV-stained glycolytic (CD45.2^+^) or OXPHOS (CD45.1^+^) T_H_17s were equally mixed and transferred into C57Bl/6 mice. (F) MitoTracker Green (MTG), (G) BCL-XL, (H) BIM, or (I, J) SCENITH metabolic phenotype were assessed in CTV^+^ vs. CD45.1^+^ live-gated cells (n=3 mice). All data are mean ±SEM. Data represent one of two independent experiments. (A-J) Statistical analysis was determined by two-way ANOVA (*p<0.05; **p<0.01; ***p<0.001; ****p<0.0001).

Next, to distinguish the relative contribution of cell death and homeostatic proliferation to the persistence of transferred cells, congenically distinct glycolytic and OXPHOS T_H_17s were generated, stained with CTV, mixed in 1:1 proportion, and adoptively transferred into lymphoreplete recipient mice. Between day 2 and 5 post co-transfer, we observed an enhanced contraction of cells that were polarized under glycolytic conditions (**Fig. 6C**). By day 5 post co-transfer, cells that originated from OXPHOS conditions exhibited better survival, less cell death, and more proliferation than cells that were polarized under glycolytic conditions (**Fig. 6C-E, S4D**). Together, these data suggest that T_H_17s polarized under OXPHOS conditions persist better than their glycolytic counterparts due to their capacity to proliferate and resist cell death.

To examine how distinct mitochondrial programs established by the use of glycolysis and OXPHOS affects T_H_17 persistence in vivo, we performed co-transfer experiments, as in (**Fig. 6C**), and characterized cells that escaped the contraction phase. Despite prominent differences in mitochondrial content and BCL-XL expression between glycolytic and OXPHOS T_H_17s at the time of co-transfer, recovered cells had similar mitochondrial mass and BCL-XL expression (**Fig. 6F, 6G**). In contrast, BIM expression remained low in the OXPHOS cultures (**Fig. 6H**). Together, this suggested that the cells with more mitochondrial content, higher BCL-XL and lower BIM expression had greater survival potential.

In parallel, we also assessed if the metabolic phenotype of in vitro T_H_17s was maintained under physiologic conditions. To do this, we utilized SCENITH, a flow-cytometry based platform that measures protein translation as a surrogate readout of metabolic activity (Arguello et al., 2020). Metabolic phenotype is inferred by the ability to translate protein following inhibition of glycolysis or OXPHOS. Importantly, SCENITH has been adapted for rare immune populations and allows for extremely rapid metabolic profiling *ex vivo*, a limitation with other methods that assess metabolism (Binek et al., 2019; Llufrio et al., 2018).

Analysis of T_H_17 cultures prior to adoptive transfer (day 0) reproduced our findings using seahorse flux analysis (**Fig. 1**). Namely, in vitro OXPHOS T_H_17s exhibited maximal mitochondrial dependency and no glucose capacity; whereas, glycolytic T_H_17s had greater glucose capacity and lower mitochondrial dependency (see day 0: **Fig. 6I, 6J, S4E**). Following co-transfer of glycolytic and OXPHOS T_H_17s into recipient mice, parallel analysis on recovered cells revealed that the surviving cells now exhibited a similar metabolic phenotype; i.e., heightened dependency on mitochondrial metabolism and low glycolytic capacity (see day 5: **Fig. 6I, 6J, S4F**). In sum, these in vivo data illustrate that OXPHOS T_H_17s maintain their metabolic and survival phenotype in vivo. In contrast, glycolytic T_H_17s exhibit a metabolic disadvantage, whereby they adapt or are selected to be more dependent on mitochondrial metabolism, the preferred in vivo metabolic phenotype, in order to survive.

### Adoptive transfer of OXPHOS T_H_17s promotes anti-tumor immunity

In preclinical studies of adoptive T cell transfer-based immunotherapy (ACT), transfer of tumor-specific T_H_17s enhances anti-tumor immunity due, in part, to their ability to persist long-term (Kryczek et al., 2011; Martin-Orozco et al., 2009; Muranski et al., 2008; Muranski et al., 2011). To examine whether OXPHOS activity during T_H_17 polarization facilitates anti-tumor immunity by supporting persistence in vivo, we adoptively transferred tumor-specific glycolytic or OXPHOS T_H_17s into tumor-bearing mice. In brief, we subcutaneously injected congenically distinct mice with Yumm5.2 melanoma cells that stably expressed the ovalbumin antigen (OVA). We employed OT-II T cell receptor transgenic mice as the source of naïve CD4 T cells to generate OVA-specific, glycolytic and OXPHOS T_H_17s, which were individually transferred into tumor-bearing mice. Tumors in mice that received OXPHOS T_H_17 cultures grew slower than those that received glycolytic T_H_17 cultures (**Fig. 7A**), and differences in tumor growth correlated with the abundance of transferred cells in the tumor draining lymph node (TDLN) and tumor (**Fig. 7B**). Analysis of the tumor microenvironment (TME) identified that the transferred cells were similar in their identity and functionality (**Fig. 7C**); however, transfer of OXPHOS T_H_17s led to more IFNγ-expressing CD8 T cells and less immunosuppressive Tregs (**Fig. 7D, 7E**). This indicated that transferring cells with a greater capacity to persist can modulate the composition and effector function of tumor infiltrating T cells. In all, these data illustrate that the metabolic phenotype of T_H_17s regulates their persistence in vivo, which helps to sustain the anti-tumor immune response in the TME.

**Fig 7:**
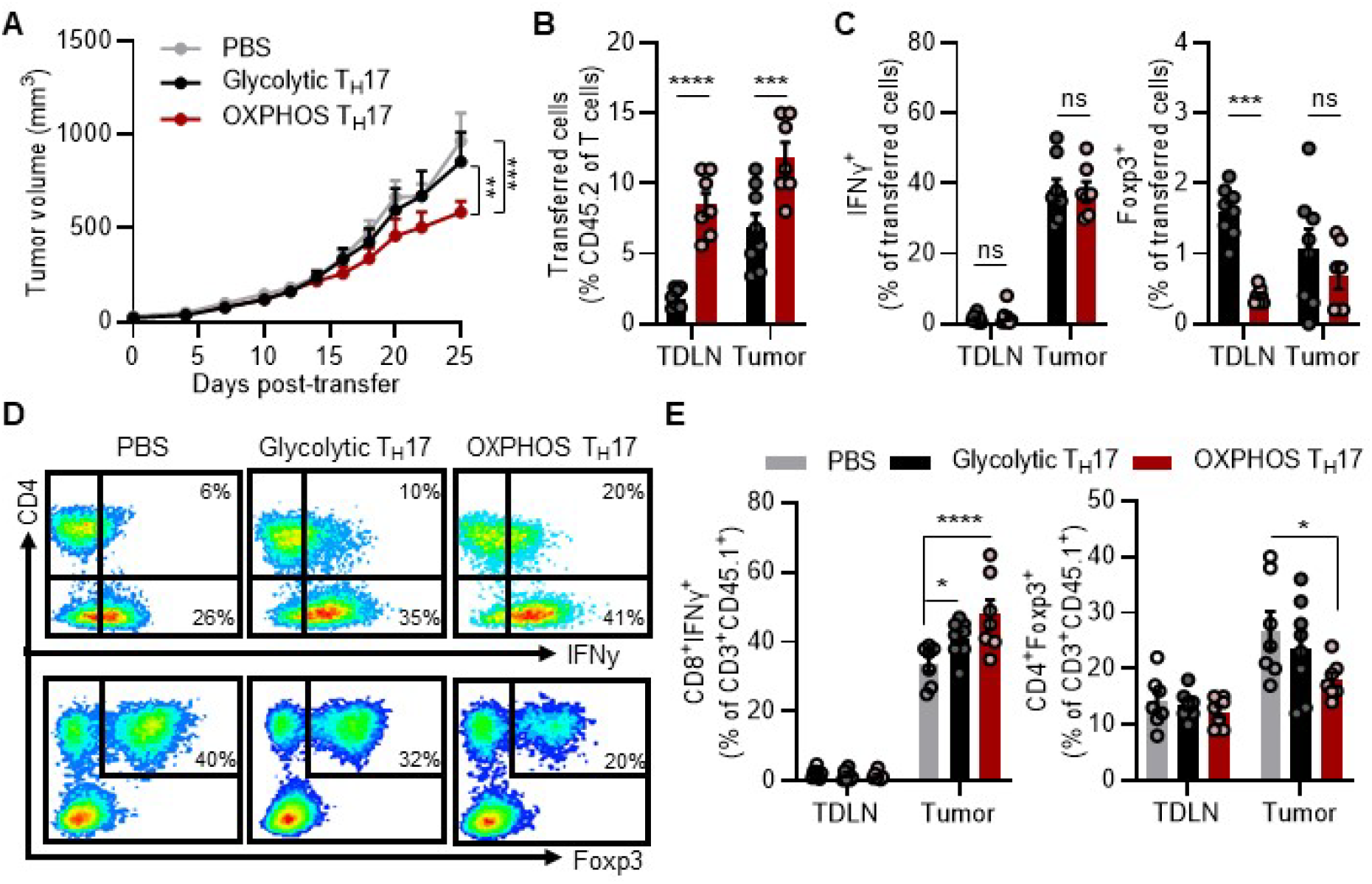
T_H_17 persistence promotes anti-tumor immunity in vivo. (A) Yumm5.2-OVA tumor growth in CD45.1 mice following transfer of glycolytic or OXPHOS OT-II CD45.2 T_H_17s (PBS control n=10; Glycolytic T_H_17s n=9; OXPHOS T_H_17s n=8). (B) Frequency and (C) phenotype of transferred cells in tumor draining lymph node (TDLN) or tumor. (D and E) Characterization of host (CD45.1^+^) T cells in the TDLN or tumor. (D) Representative flow plots and (E) accompanying quantification of IFNγ^+^ or Foxp3^+^ expression in CD45.1^+^ T cells. All data are mean ±SEM. Data represent one of two independent experiments. Statistical analysis was determined by two-way ANOVA (*p<0.05; **p<0.01; ***p<0.001; ****p<0.0001).

## Discussion

Naturally arising T_H_17s require OXPHOS, a metabolic dependency that is not readily observed in vitro and, thus, has limited our understanding of T_H_17 metabolism. In the present study, we compared T_H_17s that predominately used glycolysis or OXPHOS and demonstrated that OXPHOS activity induces an anti-apoptotic program in T_H_17s, resulting in greater persistence and enhanced anti-tumor immunity. We propose a metabolic link that regulates T_H_17 apoptotic resistance and provide insight into T_H_17 survival and its regulation by mitochondrial integrity and activity.

Herein, we first found that aerobic glycolysis is not required for T_H_17 differentiation. Seahorse flux analysis, metabolomics, and single-cell analysis using the flow cytometry-based platform SCENITH revealed our in vitro OXPHOS culture system produced cells that had little capacity to use or upregulate glycolysis and, thus, were largely reliant on OXPHOS. Under this backdrop, we found that OXPHOS can support T_H_17 differentiation and effector function to a similar degree as cells that predominately use glycolysis. These results closely align with studies demonstrating that T_H_17s rely on OXPHOS for early signaling events during differentiation and for effector function in vivo (Franchi et al., 2017; Kaufmann et al., 2019; Omenetti et al., 2019; Shin et al., 2020). Additionally, our findings do not negate the use of glycolysis in T_H_17s, instead they demonstrate that OXPHOS has the capacity to support T_H_17 lineage commitment. Although, the metabolic and signaling cues that promote the predominate use of OXPHOS in T_H_17s in vivo remain unresolved.

Second, we demonstrate that OXPHOS activity in T_H_17s promotes apoptotic resistance. Molecular analysis of BCL-2 family proteins and cell death assays confirmed distinct apoptotic thresholds in glycolytic and OXPHOS T_H_17s. Chemical inhibition of BCL-2 family proteins and, conversely, retroviral induction of BCL-XL and BIM in glycolytic and OXPHOS T_H_17s, respectively, normalized apoptotic cell death, validating the role of OXPHOS in T_H_17 apoptotic resistance. Indeed, prior work has demonstrated a link between glycolysis and peripheral deletion of T cells, establishing a model that loss of bioenergetic activity leads to mitochondrial dysfunction and precedes commitment to apoptotic cell death (Rathmell et al., 2000).

Third, we found that the anti-apoptotic program mediated by OXPHOS has lasting effects on T_H_17 cell contraction and longevity in vivo. Through adoptive transfer experiments, we show that OXPHOS T_H_17s with more mitochondria, higher levels of BCL-XL, and lower levels of BIM exhibited better persistence in vivo, relative to their glycolytic counterparts. This is in agreement with prior observations that T cells from BIM-/-mice or mice that constitutively express BCL-XL have defects in cell death that lead to an accumulation in cell number and autoimmunity (Bouillet et al., 1999; Grillot et al., 1995; Issazadeh et al., 2000).

Additionally, this study adds to the mounting number of reports illustrating that OXPHOS activity promotes the generation and maintenance of memory T cells. In the context of ACT immunotherapy, T_H_17s mediate tumor regression through indirect and direct mechanisms (Knochelmann et al., 2020; Martin-Orozco et al., 2009; Muranski et al., 2008; Muranski et al., 2011). We illustrate that OXPHOS T_H_17s exhibit the desired features of CD8 T cells that improve anti-tumor activity, namely low oxidative stress, high proliferative potential, and enhanced persistence in vivo. Although, differences in tumor growth were not due to differences in IFNγ or Foxp3 expression in transferred cells in the TME. Our findings suggest that differences in the frequency of transferred cells, or in the quality of persisting T cells present in the TME, modulate tumor growth; however, the mechanisms by which they regulate the type and functionality of tumor-infiltrating T cells warrants further investigation.

Furthermore, T_H_17s are thought to resemble memory CD8 T cells, as they exhibit similar metabolic phenotypes and long-lived capabilities. It remains to be determined whether the anti-apoptotic profile mediated by OXPHOS is specific to T_H_17s or a general phenomenon of T cells that predominately rely on OXPHOS. This knowledge will be of interest for ACT immunotherapy. The efficacy of which is dependent on the cytotoxic function and persistence of infused T cells, qualities established during in vitro expansion and determined by their cellular and metabolic state prior to transfer (Chan et al., 2021; Chang and Pearce, 2016; Kishton et al., 2017; Sukumar et al., 2016). Our data demonstrate that OXPHOS polarization produces metabolically fit T_H_17s, and that these subsequently exhibit a superior survival advantage compared to that of glycolytic cultures.

Lastly, perhaps most importantly, our study highlights the need to study T_H_17s in their native metabolic state. Recent technological advances have set the stage for in vivo and ex vivo metabolic profiling in immune cells (Voss et al., 2021). For instance, SCENITH has increased the feasibility and depth of immunometabolism studies by employing a flow cytometry-based readout, enabling analysis of sparce immune cell populations (Lopes et al., 2021). In vivo isotope tracing metabolomics is an area of active development that will enhance our understanding of immunometabolism in situ (Glick et al., 2014; Ma et al., 2019). However, there remains considerable challenges to studying metabolism in CD4 T cell subsets in vivo, as they localize to tissues that require lengthy digestion protocols and lack unique surface receptors that would otherwise enable their rapid isolation necessary for metabolic studies. Thus, in order to comprehensively examine metabolism in T_H_17s, we modeled their in vivo metabolic activity in T_H_17s generated in culture. We confirmed that our system produced OXPHOS T_H_17s that mirrored the metabolic activity of in vivo T_H_17s and validated that our OXPHOS culture system induces a metabolite profile previously shown to be critical for T_H_17 identity. We provide evidence that a physiological role of OXPHOS in T_H_17s is to control the apoptotic threshold in favor of cellular persistence in vivo, and thus may be leveraged in the setting of ACT or T_H_17 pathology.

## Acknowledgements

The authors would like to thank Dr. Kathryn V. Tormos at Agilent for helpful resources and experimental support. HSH was supported by 2T32AI007413 and T32DK094775. CAL was supported the NCI (R37CA237421) and UMCCC Core Grant (P30CA046592). Metabolomics studies performed at the University of Michigan were supported by NIH grant DK097153, the Charles Woodson Research Fund, and the UM Pediatric Brain Tumor Initiative.

## Author Contributions

HSH, AWO, LF, CAL conceived of and designed this study. HSH, LF, and CAL guided the research and wrote the manuscript. HSH, NEM, MS, KL, LL, PS, AA, AH, BM, ZL, YX, LZ, YLL, RJA, and IK provided key reagents, performed experiments, and/or analyzed data. LF, NK, IK, WZ, and CAL provided expertise in experimental design and data interpretation. CAL supervised the work carried out in this study.

## Materials and Methods

### Mice

All animal studies were performed in accordance with the Institutional Animal Care and Use Committee at the University of Michigan. All mice were on a C57BL/6 background, used at age 6-8 weeks, and maintained under specific pathogen-free housing. Wild type C57BL/6, B6.SJL (CD45.1), B6.Cg-Tg(TcraTcrb)425Cbn/J (OT-II), and RAG2^−/−^ mice were obtained from the Jackson Laboratories.

### In vitro T cell culture

Naïve CD4 T cells were isolated from mice spleens and lymph nodes by magnetic bead separation (Miltenyi Biotec) following the manufacturers’ protocols. Cells were cultured in glucose-free RPMI (Gibco) supplemented with 10% heat-inactivated FBS (Corning), 1% penicillin/streptomycin (Gibco), 50μM 2-Mercaptoethanol (Gibco), and either 10mM glucose (Sigma) or 10mM galactose (Sigma). Naïve CD4 T cells were activated with plate-bound anti-CD3e (5μg/ml; clone 145-2C11; eBioscience) and soluble anti-CD28 (2μg/ml; clone 37.51; eBioscience) in the presence of mIL-1β (10 ng/ml; R&D Systems), mIL-23 (10ng/ml; R&D Systems), mIL-6 (50ng/ml; R&D Systems), and hTGF-β (5ng/ml; Peprotech) for T_H_17 cell polarization; mIL-12 (10ng/ml; R&D Systems) for T_H_1 cell polarization, or TGF-β (10ng/ml; Peprotech) and mIL-2 (10ng/ml; R&D Systems) for T_reg_ polarization. All cells were cultured at 37°C and 5% CO_2_. T_H_17 differentiation was verified on day 3 or 4, and cytokine expression on day 5, at which point cells were used for subsequent assays.

### Flow cytometry

For live cell staining, T_H_17s (1 × 10^6^ cells/ml) were washed and incubated with metabolic dyes in serum-free media for 20 minutes at 37°C, followed by antibody surface staining in fluorescence-activated cell sorting (FACS) buffer (PBS +2% HI FBS and 2mM EDTA). For apoptosis detection, T_H_17s (1 × 10^6^ cells/mL) were plated in triplicate into a 96-well plate and treated with a cell death stimulus for 6-24 hours. Cells were washed and resuspended in 1X Annexin Binding Buffer (Thermo). Annexin V was added and allowed to incubate for 20 minutes at room temperature in the dark. Cells were washed, resuspended in 7-AAD staining solution, and immediately analyzed.

For intracellular staining, cells were surface stained, fixed and permeabilized using Foxp3/Transcription Factor Fixation/Permeabilization Concentrate and Diluent (eBioscience), and then intracellular stained. For cytokine staining, cells were stimulated with PMA (10ng/mL; Sigma-Aldrich) and ionomycin (0.5μg/mL; Sigma-Aldrich) in the presence of brefeldin A (1:1,000; eBioscience) and monensin (1:1,000; BD Biosciences) for 4 hours prior to intracellular staining.

For co-transfer experiments, peripheral (axillary, brachial, inguinal) lymph nodes and spleens were homogenized and passed through a 40μm cell strainer to create single cell suspensions. Splenocytes and blood samples were incubated in red blood cell lysis buffer (eBioscience) for 5 minutes at room temperature, washed in PBS, and resuspended in FACS buffer prior to staining. For isolation of lamina propria lymphocytes, harvested colons were cut longitudinally, washed in PBS, dissected into small pieces, washed in HBSS media supplemented with 2.5% HI FBS and 1% penicillin/streptomycin, and then incubated in the presence of 1mM DTT for 15 minutes at 37°C with 200rpm shaking to remove mucus. Then intestinal pieces were incubated in supplemented HBSS media containing 1mM EDTA for 30 minutes at 37°C with 200rpm shaking, washed, and 1mM EDTA incubation repeated, followed by further digestion with 1mM collagenase type III (Sigma) and DNase for 2 hours. Digested tissues were washed, filtered, resuspended in 40% Percoll, layered onto 75% Percoll, and centrifuged for 20 minutes at 2,000rpm. Lamina propria lymphocytes in the interphase were collected and washed prior to flow staining.

For tumor in vivo experiments, single cell suspensions were prepared from mouse tumor tissue and tumor draining lymph node (TDLN). Tumor infiltrating lymphocytes were enriched by density gradient centrifugation prior to intracellular cytokine staining, as detailed above.

The following antibodies were from BD Biosciences, eBioscience, or Biolegend: anti-mouse CD90 (clone 53-2.1), anti-mouse CD45.1 (clone A20), anti-mouse CD45.2 (clone 104), anti-mouse TCRβ (clone H57-597), anti-mouse CD4 (clone GK1.5 and RM4-5), anti-mouse CD8 (clone 53-6.7), anti-mouse RORγt (clone AFKJS-9), anti-mouse Foxp3 (clone FJK-16s), anti-mouse Tbet (clone 4B10), anti-mouse IL-17A (clone TC11-18H10), anti-mouse IFN-γ (clone XMG1.2), anti-Bcl-2 (clone BCL/10C4), and goat anti-Rabbit IgG (H+L) Highly Cross-Absorbed Secondary Antibody (2μg/mL; Invitrogen). Annexin V (Invitrogen), 7-AAD (Invitrogen), Live/dead fixable viability dye (Invitrogen), H_2_DCFDA (1μM; Invitrogen), TMRM (25nM; Invitrogen), MitoSOX Red Mitochondrial Superoxide Indicator (5μM; Invitrogen), MitoTracker Green FM (20nM; Invitrogen), and CellTrace Violet Cell Proliferation (5μM; Invitrogen) were purchased from ThermoFisher Scientific. Anti-Bcl-xL (54H6; 1:800) and anti-Bim (C34C5; 1:400) were purchased from Cell Signaling Technology. Flow samples were acquired using a Fortessa analyzer (BD Biosciences), MoFlow Cell Sorter (Beckman Coulter), or ZE5 Cell Analyzer (Bio-Rad). Data were analyzed with DIVA software (BD Biosciences) or FlowJo software (TreeStar).

### Seahorse

Seahorse assays were performed using a XF-96 Extracellular Flux Analyzer (Agilent). Sensor cartridges were incubated in dH_2_O overnight, and then hydrated in XF calibrant (Agilent) for 1 hour in a non-CO_2_ incubator at 37°C on the day of the assay. Hydrated cartridges were loaded with oligomycin (1μM), FCCP (1μM), rotenone (0.1μM), and antimycin A (1μM) for the mitostress test; or rotenone (0.5μM), antimycin A (0.5μM), and 2-deoxy-glucose (50mM) for the glycolytic rate assay. Differentiated T_H_17s were washed and resuspended in Seahorse XF RPMI media (Agilent;103576) supplemented with XF Glutamine (Agilent;103579) and either 10mM glucose or 10mM galactose. 300,000 cells per well were seeded on poly-L-lysine-coated plates and allowed to equilibrate for 30 minutes in a non-CO_2_ incubator at 37°C. After the assay, measurements were normalized based on cell seeding density using CyQuant (Invitrogen). For the mitostress test, the metabolic phenotype was determined based on basal OCR and ECAR measurements (i.e., prior to inhibitor treatment). SRC was determined by subtracting basal OCR from maximal OCR measurements. For the glycolytic rate assay, PER measurements were calculated by the GRA Report Generator using Wave 2.3 software.

### SCENITH

SCENITH was performed as described in (Arguello et al., 2020). SCENITH reagents kit (inhibitors, puromycin, and antibodies) were obtained from www.scenith.com/try-it and used according to the provided protocol for T lymphocytes. In brief, in vitro T_H_17s were resuspended in RPMI containing 10% HI FBS and either 10mM glucose or 10mM galactose and plated in a 96-well plate at 1×10^6^ cells/mL in triplicate. Cells were pretreated with 2-deoxy-glucose (50mM), oligomycin (1μM), or in combination for 15 minutes, followed by treatment with puromycin (10μg/mL) for 30 minutes at 37°C. After drug treatments, samples were placed on ice and cells were washed with cold PBS and surface stained for 15 minutes at 4°C in FACS buffer. After fixation and permeabilization, cells were intracellular stained with anti-puromycin (1:500; clone R4743L-E8) for 1 hour at 4°C. For ex vivo studies, harvested lymph nodes were immediately processed to generate single-cell suspensions in prewarmed RPMI containing 10% HI FBS and 10mM glucose in FACS tubes and treated as described above.

### Metabolomics

Metabolites were extracted from T_H_17s by adding cold 80% methanol, incubating at - 80°C for 10 minutes, followed by centrifugation at 10,000xg for 10 minutes at 4°C. Supernatants were collected and lyophilized by speedvac. The quantity of metabolite fraction analyzed was normalized to the corresponding protein concentration calculated upon processing a parallel replicate. For isotope tracing metabolomics of in vitro cells, T_H_17s were cultured in glutamine-free RPMI supplemented with equimolar concentrations of unlabeled glutamine or [U]13C-glutamine (Cambridge) overnight prior to metabolite extraction. Alternatively, glycolytic T_H_17s were cultured in glucose-free RPMI supplemented with 10mM unlabeled glucose or [U]13C-glucose (Cambridge). In parallel, OXPHOS T_H_17s were cultured in glucose-free RPMI supplemented with 10mM unlabeled galactose or [U]13C-galactose (Cambridge) prior to metabolite extraction. For [U]13C-glutamine tracing of T_H_17 cells generated in vivo vs in vitro, T_H_17s were activated and purified as previously described (Franchi et al., 2017), and cultured for 3 hours in RPMI containing either unlabeled glutamine or [U]13C-glutamine. Metabolite extracts were normalized to cell number. Liquid chromatography-based targeted tandem mass spectrometry (LC-MS/MS)-based metabolomics were performed and the data analyzed as previously described (Halbrook et al., 2019; Lee et al., 2019; Yuan et al., 2019).

### Immunoblotting

Cells were lysed in cell lysis buffer supplemented with protease and phosphatase inhibitors, and protein concentrations were determined with BCA Protein Assay (Pierce). Proteins were separated by SDS-PAGE followed by immunoblotting with the following Cell Signaling antibodies: anti-Bcl-2 (3498), anti-Bcl-xL (2764), anti-Mcl-1 (5453), anti-Bim (2933), anti-Bib (2003), anti-Bak (12105), anti-caspase 3 (9662), anti-cleaved caspase 3 (9664), anti-PARP (9542), anti-p-DRP1 (3455), anti-DRP1 (8570), anti-OPA1 (80471), anti-vinculin (13901), anti-tubulin (2144), HRP-conjugated anti-Rabbit IgG (7074), and PCG1α (ab54481) was from Abcam.

### Viral plasmids, transfection, and transduction

pMIG (empty vector control; 9044), pMIG Bcl-XL (8790), and pMIG Bim (8786) retroviral constructs were purchased from Addgene. Retroviruses were produced by the Vector Core at the University of Michigan. Naïve CD4 T cells were activated under glycolytic or OXPHOS conditions and in the presence of T_H_17 polarizing cytokines for 48 hours. Retroviral supernatants were loaded by centrifugation (2,000xg for 2 hours at 30°C) onto non-tissue culture treated 6-well plates pre-coated with RetroNectin (20μg/mL; Takara). Activated T cells were added and spin-transduced for 30 minutes at 1,000xg, 30°C. Cells were expanded and transduced GFP^+^ cells were gated and used for analysis.

Parental Yumm5.2 melanoma cells were transduced with lentivirus containing pLVX-puro-OVA. Lentivirus was produced by transfecting 293FT cells and viral supernatant was collected and passed through a 0.45um filter. Following transduction, stable cell lines were established post-puromycin selection. Cell lines were cultured in DMEM F-12 media (Gibco), supplemented with 10% FBS (Corning), HEPES (Gibco), and 1% NEAA (Gibco), and regularly tested for mycoplasma contamination using MycoAlert (Lonza). pLVX-puro-OVA (Addgene; 135073) and Yumm5.2 cells were generous gifts from Dr. Maria Castro and Drs. Ilona Kryczek and Weiping Zou, respectively.

### In vivo mouse experiments

For all transfer experiments, donor and recipient mice were age- and sex-matched. For persistence studies, CD45.2^+^ glycolytic and CD45.1^+^ OXPHOS T_H_17s were PBS washed, mixed in equal proportions, and a total of 10^6^ T_H_17s were transferred into RAG2^−/−^ mice through tail vein injection. For competition studies, CD45.2^+^ glycolytic and CD45.1^+^ OXPHOS T_H_17s were stained with Cell Trace Violet, washed with PBS, mixed in equal proportions, and then a total of 4×10^6^ T_H_17s were transferred into C57BL/6 mice via tail vein injection. For tumor studies, CD45.1 mice were s.c. injected with 75,000 Yumm5.2-OVA melanoma cells prepared in a 1:1 mixture of growth factor reduced matrigel (Corning) and PBS. 10^7^ OT-II (CD45.2^+^) glycolytic or OXPHOS T_H_17s were transferred via tail vein injection. Tumors were measured every 3 days using digital calipers and tumor volumes were calculated based on the formula (length x width x width)/2.

### RNAseq

Total RNA was extracted from Glycolytic and OXPHOS T_H_17s using RNeasy Plus Micro Kit (Qiagen). Strand-specific, polyA selected libraries were prepared and paired-end, 100-base reads were sequenced using NovaSeq-6000 (Illumina).

### Statistical analysis

The data are shown as means ±SEM. For flow cytometry analysis, median fluorescent intensity was determined by FlowJo software and used for analyses. Statistical analyses were performed using GraphPad Prism software and statistical significance was determined using the following analyses: paired t-tests (to compare two matched experimental groups from pooled independent experiments), unpaired t-test (to compare two matched experiment groups performed in technical triplicates), or one-way ANOVA (to compare one factor among three groups or more) and two-way ANOVA (to compare two factors among three groups or more) using Turkey’s multiple comparisons test. P-value<0.05 was considered significant, unless noted otherwise. Type and number of experimental replicates and explanation of significant values are presented within the legends.

## Supplemental Figures

**Supplemental Figure 1:**
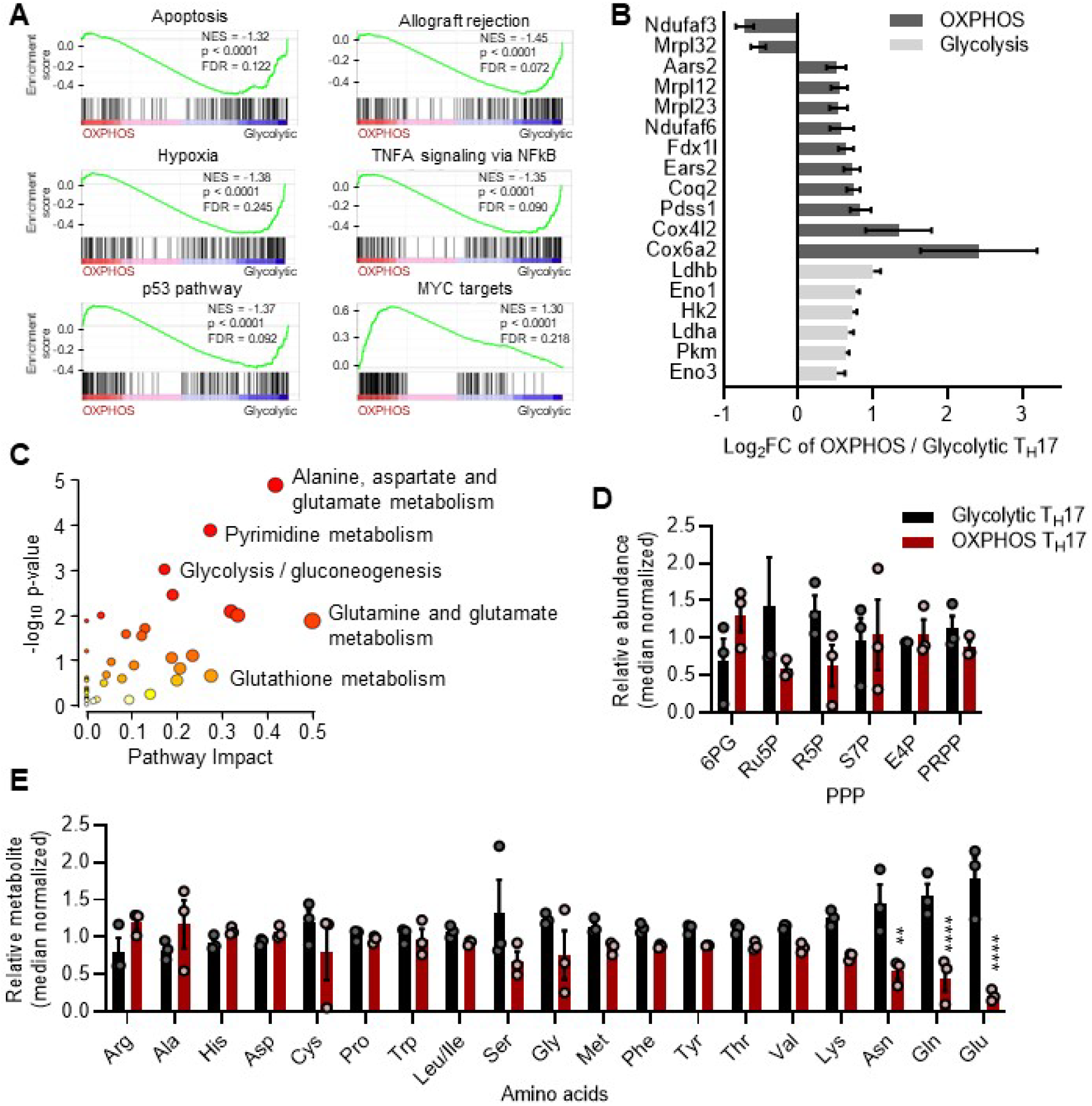
Transcriptomic and metabolic interrogation of OXPHOS T_H_17s. (A) Gene set enrichment or (B) expression of metabolic genes (log_2_FC >|0.5|, p-value<0.05) in OXPHOS vs. glycolytic T_H_17s analyzed by RNAseq. (C) Pathway enrichment of altered metabolites (log_2_FC >|1|, p-value<0.1). (D) Relative abundances of pentose phosphate pathway metabolites (PPP) and amino acids measured by metabolomics. All data are mean ±SEM (n=3 biological replicates). (D,E) Statistical analysis was determined by two-way ANOVA (*p<0.05; **p<0.01; ****p<0.0001).

**Supplemental Figure 2:**
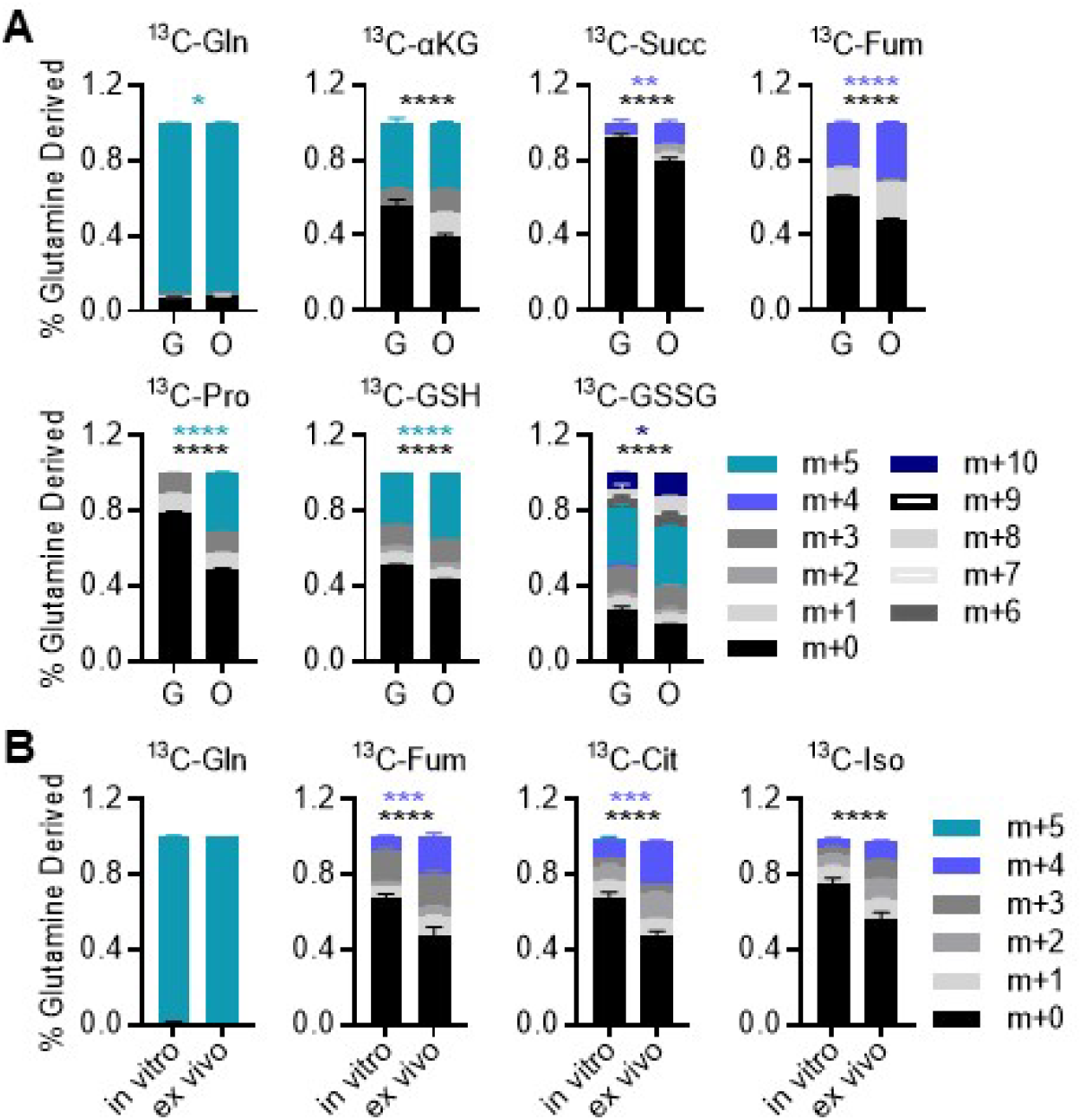
Glutamine metabolism in OXPHOS T_H_17s. (A and B) [U]^13^C glutamine tracing metabolomics in (A) glycolytic vs. OXPHOS T_H_17s or (B) T_H_17s generated in vitro vs. in vivo. Fractional labeling pattern of isotopologues for indicated metabolites (mass+0-10; m+): ^13^C glutamine (^13^C-Gln; m+5), ^13^C glutamine-derived α-ketoglutarate (^13^C-αKG; m+5), succinate (^13^C-Succ; m+4), fumarate (^13^C-Fum; m+4), proline (^13^C-Pro; m+5), reduced glutathione (^13^C-GSH; m+5), and oxidized glutathione (^13^C-GSH; m+5, m+10), citrate (^13^C-Cit; m+4), and isocitrate (^13^C-Iso; m+4). All data represent 2-3 biological replicates, graphs depict mean ±SEM, and statistical analysis was determined by two-way ANOVA (*p<0.05; **p<0.01; ***p<0.001; ****p<0.0001).

**Supplemental Figure 3:**
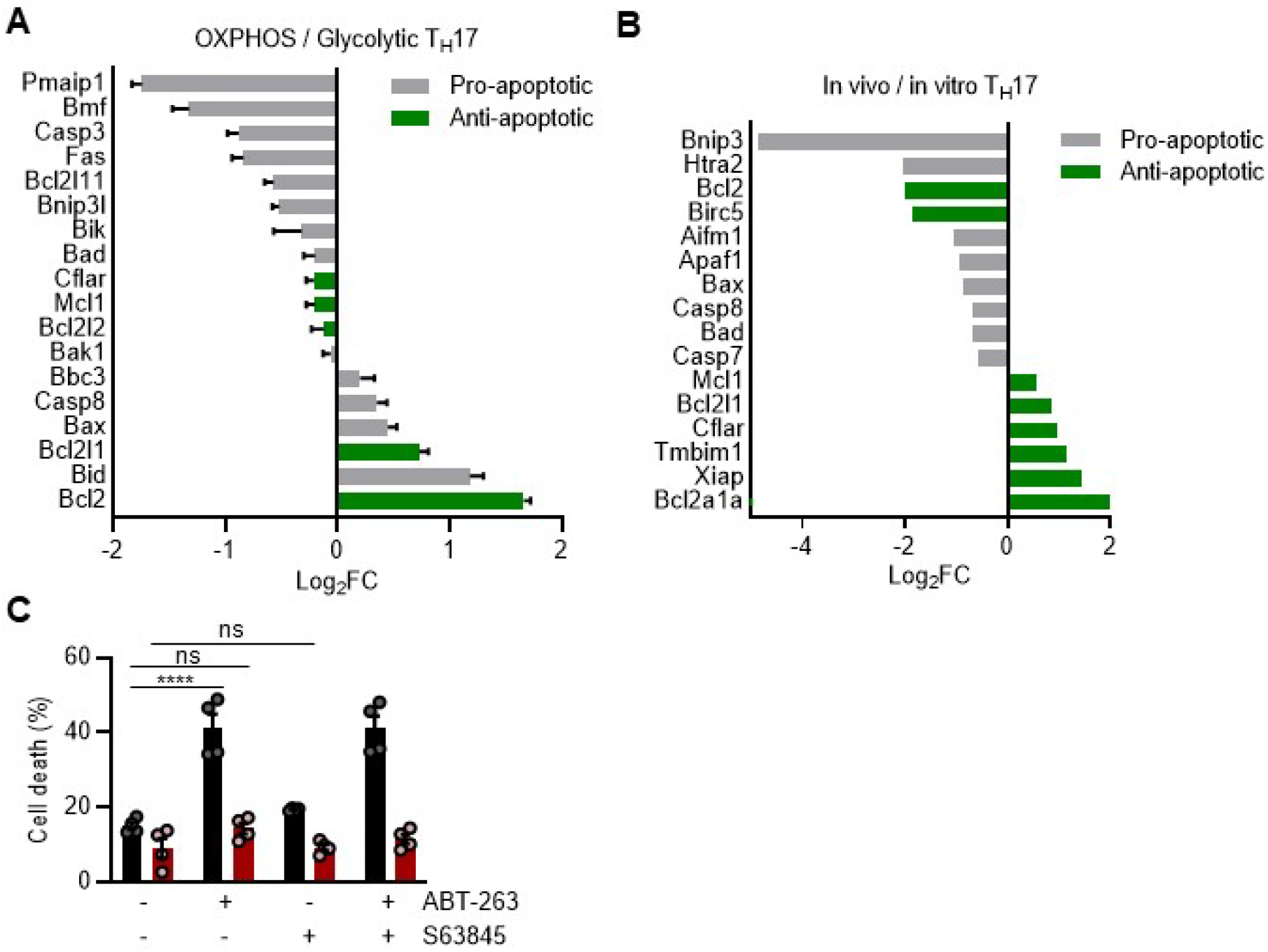
Profile of apoptotic regulator expression and functionality in T_H_17s. (A and B) Transcriptomic analysis of apoptotic regulators in (A) OXPHOS vs. glycolytic T_H_17s or (B) T_H_17s generated in vivo vs. in vitro (log_2_FC >|0.5|, p-value<0.05) (n=2-3 biological replicates). (C) T_H_17s treated with ABT-263 and/or S63845 for 24 hours, and then stained with Annexin V and 7-AAD to quantify the percentage of cell death by flow cytometry (n=4). Data represent two independent experiments. All data are mean ±SEM. (C) Statistical analysis was determined by two-way ANOVA (****p<0.0001).

**Supplemental Figure 4:**
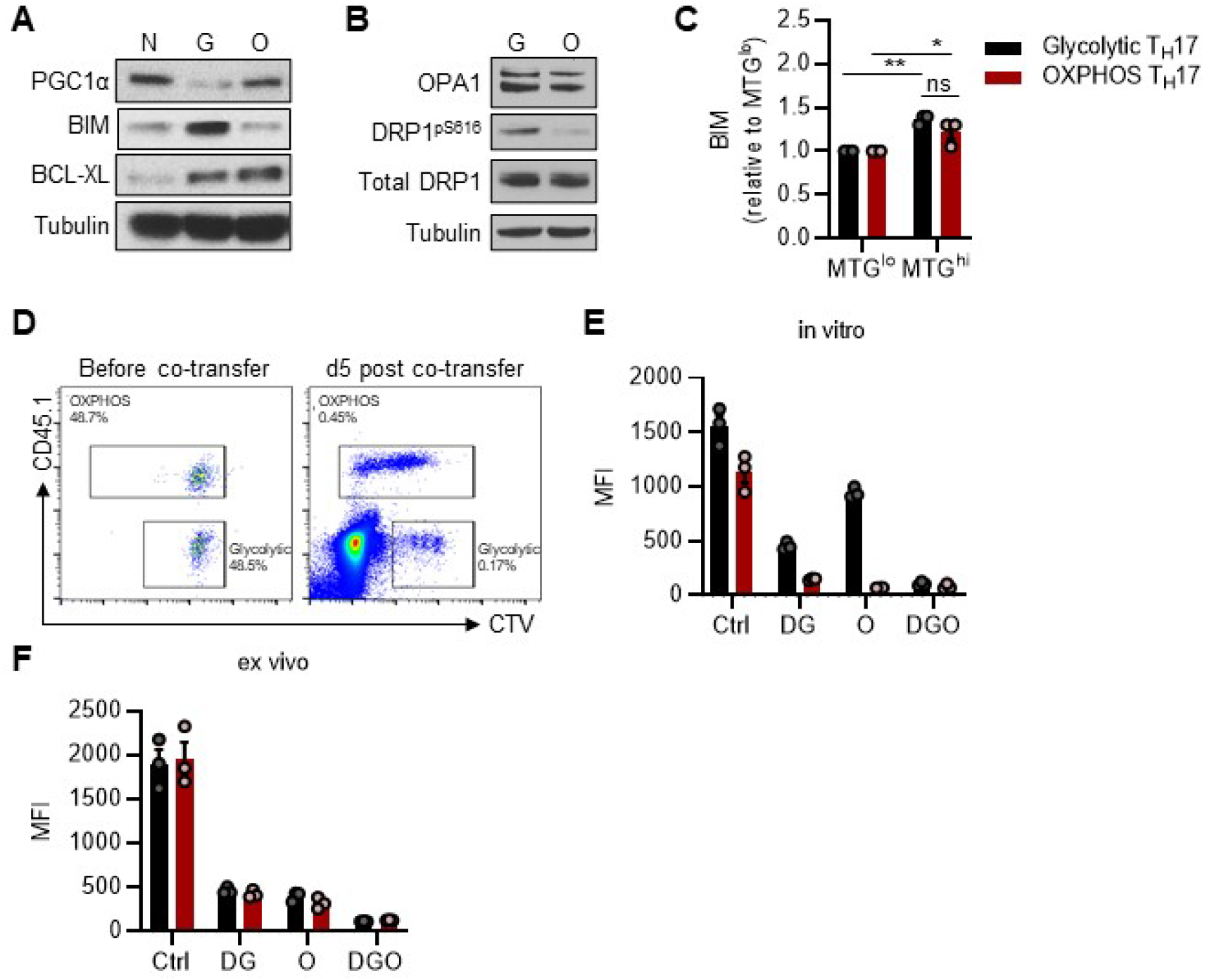
Mitochondrial integrity is critical for T_H_17 cell survival. (A and B) Representative western blots of (A) PGC1α, BIM, and BCL-XL or (B) mitochondrial fusion regulator OPA1 and fission regulator DRP1^pS616^ in naïve CD4 T cells; N, glycolytic T_H_17s; G, and OXPHOS T_H_17s; O. (C) MTG-stained T_H_17s were flow sorted into mitochondrial low and high fractions (20% lowest and highest MFI, respectively), and then stained for BIM (n=3). Data represent 3 independent experiments. (D) Glycolytic (CD45.2^+^) T_H_17s and OXPHOS (CD45.1^+^) T_H_17s stained with Cell Trace Violet (CTV), equally mixed (1:1), and then transferred into C57Bl/6 (CD45.2^+^) recipient mice. Representative flow plots of transferred cultures on day 0 (left) and 5 days post-transfer (right). (E and F) Glycolytic and OXPHOS T_H_17s were polarized in vitro and their metabolic phenotype was assessed using SCENITH (E) on the day of co-transfer or (F) 5 days following co-transfer. All data are mean ±SEM. (C) Statistical analysis was determined by two-way ANOVA (*p<0.05; **p<0.01).

